# Cross-Platform Validation of Neurotransmitter Release Impairments in Schizophrenia Patient-Derived *NRXN1*-Mutant Neurons

**DOI:** 10.1101/2020.11.03.366617

**Authors:** ChangHui Pak, Tamas Danko, Vincent R. Mirabella, Jinzhao Wang, Xianglong Zhang, Thomas Ward, Sarah Grieder, Madhuri Vangipuram, Yu-Wen Alvin Huang, Yingfei Liu, Kang Jin, Philip Dexheimer, Eric Bardes, Alexis Mittelpunkt, Junyi Ma, Michael McLachlan, Jennifer C. Moore, Alexander E. Urban, Jeffrey L. Dage, Bradley J. Swanson, Bruce J. Aronow, Zhiping P. Pang, Douglas F. Levinson, Marius Wernig, Thomas C. Südhof

## Abstract

Heterozygous *NRXN1* deletions constitute the most prevalent currently known single-gene mutation predisposing to schizophrenia. Previous studies showed that engineered heterozygous *NRXN1* deletions impaired neurotransmitter release in human neurons, suggesting a synaptic pathophysiological mechanism. Utilizing this observation for drug discovery, however, requires confidence in its robustness and validity. Here, we describe a multi-center effort to test the generality of this pivotal observation, using independent analyses at two laboratories of patient-derived and newly engineered human neurons with heterozygous *NRXN1* deletions. We show that in neurons that were trans-differentiated from induced pluripotent stem cells derived from three *NRXN1*-deletion patients, the same impairment in neurotransmitter release was observed as in engineered *NRXN1*-deficient neurons. This impairment manifested as a decrease in spontaneous synaptic events and in evoked synaptic responses, and an alteration in synaptic paired-pulse depression. *Nrxn1*-deficient mouse neurons generated from embryonic stem cells by the same method as human neurons did not exhibit impaired neurotransmitter release, suggesting a human-specific phenotype. *NRXN1* deletions produced a reproducible increase in the levels of CASK, an intracellular *NRXN1*-binding protein, and were associated with characteristic gene expression changes. Thus, heterozygous *NRXN1* deletions robustly impair synaptic function in human neurons regardless of genetic background, enabling future drug discovery efforts.

## INTRODUCTION

Schizophrenia is a devastating brain disorder that affects millions of people worldwide and exhibits a strong genetic component. In a key discovery, deletions or duplications of larger stretches of chromosomal DNA that lead to copy number variations (CNVs) were identified two decades ago (Sebat et al., 2004; Sebat et al., 2007). CNVs occur unexpectedly frequently, are often *de novo*, and usually affect multiple genes depending on the size of the deleted or duplicated stretch of DNA. Strikingly, the biggest genetic risk for schizophrenia was identified in three unrelated CNVs, a duplication of region 16p11.2 and deletions of 22q11.2 and of 2p16.3 (Kirov, 2015; Coelewij and Curtis, 2018; Malhotra and Sebat, 2012; Marshall et al., 2017; Anh et al., 2014; Kirov et al., 2014; Rees et al., 2014). Of these CNVs, 16p11.2 and 22q11.2 CNVs affect more than 20 genes, whereas 2p16.3 CNVs impact only one or more exons of a single gene, *NRXN1*, which encodes the presynaptic cell-adhesion molecule neurexin-1 (Hu et al., 2019; Coelewij and Curtis, 2018; Kasem et al., 2018; Sudhof, 2017; Kirov, 2015; Marshall et al., 2017). *NRXN1* CNVs confer an approximately ten-fold increase in risk of schizophrenia, and additionally strongly predispose to other neuropsychiatric disorders, especially autism and Tourette syndrome (Lowther et al., 2017, Castronovo et al., 2020). Moreover, genome-wide association studies using DNA microarrays identified common changes in many other genes that predispose to schizophrenia with smaller effect sizes (SCZ working group of PGC, 2014; Parnidas et al., 2018; Fromer et al., 2014; Fromer et al., 2016; Ripke et al., 2017; Sullivan and Geschwind 2019; Flaherty et al., 2019; Hall et al., 2020). Viewed together, these studies indicate that variations in a large number of genes are linked to schizophrenia. Among these genetic variations, heterozygous exonic CNVs of *NRXN1* are rare events, but nevertheless constitute the most prevalent high-risk single-gene association at present.

Neurexins are central regulators of neural circuits that control diverse synapse properties, such as the presynaptic release probability, the postsynaptic receptor composition, and synaptic plasticity (Missler et al., 2003; Aoto et al., 2013 and 2015; Anderson et al., 2015; Chen et al., 2017, Dai et al., 2019; Luo et al., 2020). To test whether heterozygous *NRXN1* mutations might cause functional impairments in human neurons, we previously generated conditionally mutant human embryonic stem (ES) cells that enabled induction of heterozygous *NRXN1* deletions using Cre-recombinase (Pak et al., 2015). We then analyzed the effect of the deletion on the properties of neurons trans-differentiated from the conditionally mutant ES cells using forced expression of Ngn2, a method that generates a relatively homogeneous population of excitatory neurons that are also referred to as induced neuronal (iN) cells (Zhang et al., 2013). These experiments thus examined isogenic neurons without or with a heterozygous *NRXN1* loss-of-function mutation that mimicked the schizophrenia-associated 2p16.3 CNVs, enabling precise control of the genetic background. In these experiments, the heterozygous *NRXN1* deletion produced a robust but discrete impairment in neurotransmitter release without major changes in neuronal development or morphology (Pak et al., 2015). These results were exciting because they suggested that a discrete impairment in neurotransmitter release could underlie the predisposition to schizophrenia conferred by the 2p16.3 CNVs, but these experiments did not reveal whether the actual *NRXN1* mutation that is observed in schizophrenia patients induces the same synaptic impairment (Hyman, 2015).

The present project was initiated to achieve multiple overlapping aims emerging from the initial study of Pak et al., 2015. First, we aimed to validate or refute the results obtained with neurons generated from engineered conditionally mutant ES cells, but now using neurons generated from patient-derived iPS cells with *NRXN1* mutations (Fig. 1A). This goal was pursued in order to gain confidence in the disease-relevance of the observed phenotypes. Second, we wanted to test whether the observed phenotype is independent of the laboratory of analysis, i.e. whether it is sufficiently robust to be replicated at multiple sites (Fig. 1A). This goal was motivated by the observation of poor reproducibility of neuroscience research. However, we hypothesized that the lack of reproducibility is often due to variations in experimental conditions rather than true failed replication testing of the original findings and designed our studies to demonstrate robustness of the findings through replication. Third, we aimed to generate reagents that could be broadly used by the scientific community for investigating the cellular basis of neuropsychiatric disorders (Panchision, 2016). This goal was prompted by the challenges posed by the finding that many different genes appear to be linked to schizophrenia. Fourth, we aimed to definitively establish or exclude the possibility that human neurons are uniquely sensitive to a heterozygous loss of *NRXN1* as compared to mouse neurons (Fig. 1B). The goal here was to establish whether at least as regards to *NRXN1*, mouse and human neurons exhibit fundamental differences. Fifth and finally, we hoped to gain further insights into the mechanisms by which *NRXN1* mutations may predispose to schizophrenia, an obviously needed objective given our lack of understanding of this severe disorder. As described in detail below, our data provide advances towards meeting these goals, establishing unequivocally that heterozygous *NRXN1* deletions in human but not in mouse neurons cause a robust impairment in neurotransmitter release that is replicable in multiple laboratories.

**Figure 1:**
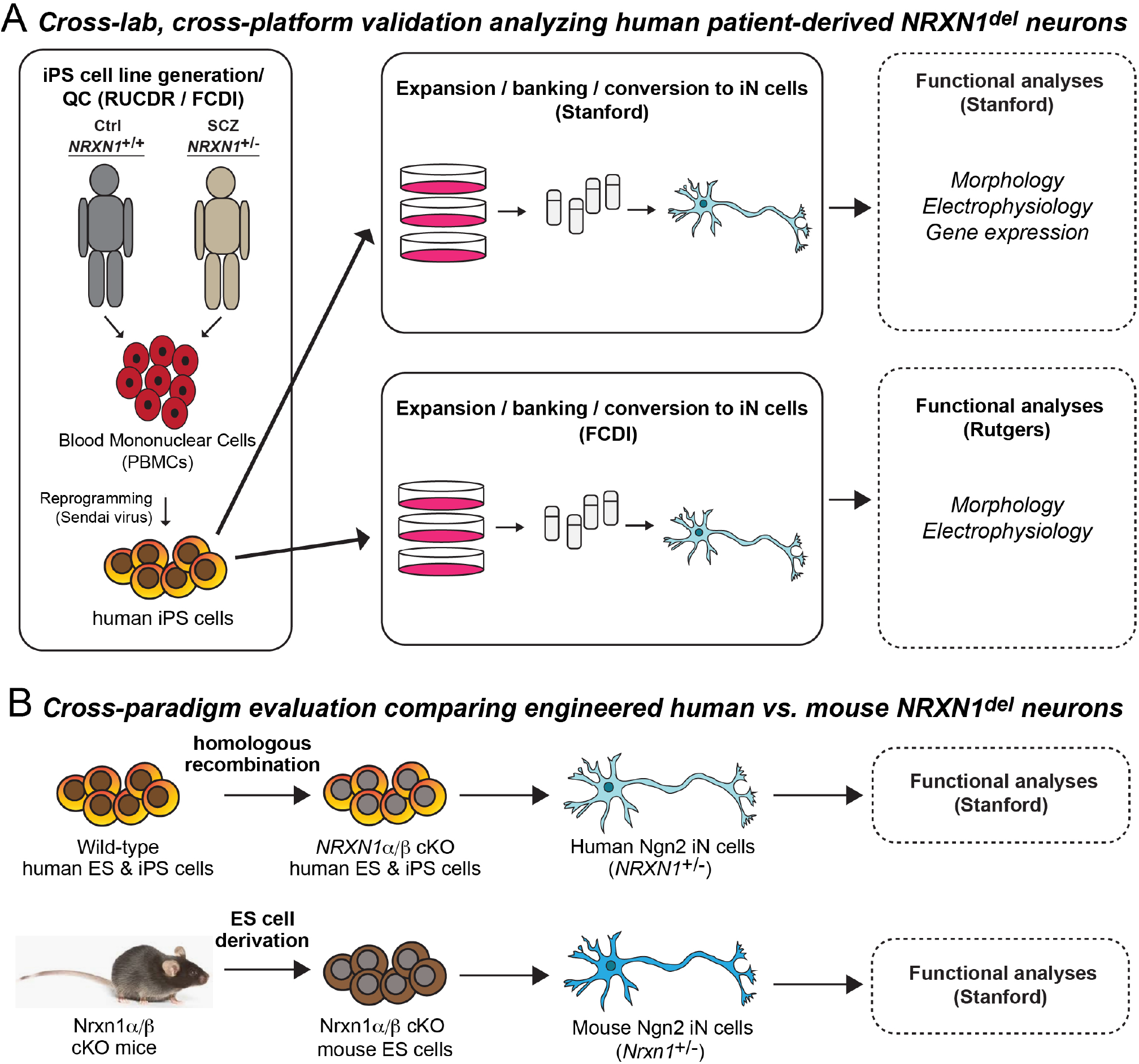
Overall study design illustrating the experimental approach to analyze human heterozygous NRXN1 loss-of-function mutations, to achieve cross-lab and cross-platform validation of observed phenotypes, and to perform cross-paradigm evaluations of these phenotypes in human and mouse neurons. **A.** Experimental strategy for analyzing the functional effects of heterozygous *NRXN1* loss-of-function mutations in human patient-derived neurons and for validating the observed phenotypes in a cross-lab and cross-platform comparison. Peripheral mononuclear blood cells (PBMCs) from schizophrenia patients with *NRXN1* deletions and from control individuals were reprogrammed into iPS cells by Rutgers University (RUCDR Infinite Biologics). iPS cells that passed quality control (QC) were shipped to Stanford and to Fujifilm-CDI Inc. (FCDI) for expansion, banking, and trans-differentiation into induced neurons. The indicated subsequent analyses were carried out at Stanford University and at Rutgers University. FCDI manufactured industry-scale human induced neurons that were shipped to Rutgers for analysis, whereas Stanford generated induced neurons at an academic single lab scale for analysis. **B.** Experimental strategy to evaluate the conservation of *NRXN1*-deletion phenotypes observed in human neurons in mouse neurons (cross-paradigm evaluation). Human and mouse stem cells that carried heterozygous engineered conditional *NRXN1/Nrxn1* deletions were trans-differentiated into neurons by Ngn2 expression and analyzed using similar approaches to ensure comparability. In this approach, isogenic human and mouse neurons without or with *NRXN1/Nrxn1* deletions were compared to test whether side-by-side analysis of human and mouse neurons prepared by indistinguishable approaches yields similar phenotypes.

## RESULTS

### Cohort of cases and controls

Peripheral blood mononuclear cell specimens and genomic DNA were obtained from the NIMH Repository and Genomics Resource (NRGR), donated by schizophrenia patients carrying heterozygous *NRXN1* exonic deletions and control individuals who were participants in the Molecular Genetics of Schizophrenia (MGS2) European-ancestry cohort (Shi et al., 2009). The controls met criteria that predicted low genetic risk for schizophrenia. Cases and controls were aged 35-51 years at collection (for details, see Methods; for information on the availability of non-identified clinical information and of biomaterials, see www.nimhgenetics.org; for patient and control properties of the cell lines reported here, see Table 1). iPS cell lines were generated from peripheral blood mononuclear cells by the Rutgers University Cell and DNA Repository (RUCDR) via integration-free Sendai virus reprogramming, and three subclones were generated from each line for analysis (Fig. 1). Reprogramming included extensive quality control measures to verify normal karyotypes, morphology, and pluripotency of iPS cells, which were also tested pre- and post-freezing for mycoplasma (Fig. S2). Whole genome sequencing of genomic DNA validated the presence of an exonic *NRXN1* deletion in each patient but none in the controls (Fig. S1). No patient or control carried other exonic CNVs known to be associated with schizophrenia or contained non-synonymous single nucleotide variants except for a variant in N1884 (chr2:50724817G>A [hg19]; chr2:50497679G>A [hg38]), which is predicted to be benign.

**Table 1.** Cases and Controls.

| NCRCG_ID | Line ID | Repository ID | Pair # | Sex | Age at donation | Diagnosis | Genotype Effect | CNV | Start (hg38) | Stop (hg38) | Size (kb) | hg19 position |
| --- | --- | --- | --- | --- | --- | --- | --- | --- | --- | --- | --- | --- |
| C3141a | Ctrl. 1 | 06C53141 | 1 | M | 41 | None | Control | - |  |  |  |  |
| C2463c | Ctrl. 2 | 06C52463 | 2 | F | 39 | None | Control | - |  |  |  |  |
| C8905a | Ctrl. 3 | 05C48905 | 3 | M | 47 | None | Control | - |  |  |  |  |
| N3320a | NRXN1<sub>del 1</sub> | 03C23320 | 1 | M | 47 | SCZ | NRXN1α deletion | 2p16.3 del | 50,899,162 | 51,081,962 | 182,800 | chr2:51,126,300-51,309,100 |
| N9540c/a<sup>1</sup> | NRXN1<sub>del 2</sub> | 04C29540 | 2 | F | 35 | SCZ | NRXN1α $\beta$ deletion | 2p16.3 del | 50,346,562 | 51,397,924 | 1,051,362 | chr2:50,573,700-51,625,062 |
| N1884a | NRXN1<sub>del 3</sub> | 05C51884 | 3 | M | 51 | SCZ | NRXN1α deletion | 2p16.3 del | 50,979,562 | 51,087,962 | 108,400 | chr2:51,206,700-51,315,100 |

<sup>1</sup>Rutgers U. used N9540a iN cells grown from FCDI while Stanford used N9540c clone for differentiation.

Of the five lines with *NRXN1* deletions (referred to as *NRXN1^del^*) and the six control lines available, we selected three pairs of cases and controls based on similarity in gender and age at donation (Table 1). Two of the lines selected had deletions affecting only NRXN1α, while the third line had a deletion ablating expression of both NRXN1α and NRXN1β (Fig. S1). All subsequent differentiations, functional assays, and data analyses were performed using this pairing system, such that each patient-derived human iPS cell line was consistently paired with its own control human iPS cell line to minimize experimental and line-to-line variability. This approach does not imply genetic similarity between the cases and controls, and the pairing is by necessity intrinsically random. Frozen vials of iPS cells were shipped for cell line expansion, banking, and iN cell trans-differentiation to generate human neurons at two sites, Stanford University and FUJIFILM Cellular Dynamics Inc. (FCDI) (Fig. 1). The neurons generated at Stanford University were then also analyzed at Stanford, whereas the neurons generated at FCDI were shipped for analysis to Rutgers University.

### Single laboratory vs. industry-scale methods of Ngn2-induced trans-differentiation of iPS cells into neurons

Forced expression of Ngn2 rapidly converts human ES or iPS cells into excitatory neurons (Zhang et al., 2013; Pak et al., 2015; Yi et al., 2016). These human neurons, also referred to as iN cells, are composed of a population of excitatory, relatively homogeneous neurons resembling cortical layer 2/3 pyramidal neurons (Zhang et al., 2013). Since production of Ngn2-induced human neurons is robust and scalable, we decided to further optimize it with two specific goals in mind: 1) to generate human neurons by trans-differentiation of iPS cells that are grown on feeder cell layers, which was necessary to minimize karyotypic abnormalities, and 2) to manufacture human neurons at an industry scale, thus enabling distribution of human neurons to multiple sites for downstream functional studies.

Guided by our published protocols (Pak et al., 2018; see Methods), we modified the iPS cell dissociation step to efficiently remove mouse feeder cells from human iPS cell colonies. We also improved the initial induction step to provide the optimal medium combination for cell survival and TetO-inducible Ngn2 expression via doxycycline. However, we observed unexpected differences between iPS cell lines. Every iPS cell line pair had to be optimized separately to achieve workable conditions for the induction, differentiation and survival of Ngn2-induced neurons (Fig. S3). Adjustments of lentivirus titers, starting cell numbers, and timing of the window of doxycycline application before puromycin selection were made for each iPS cell line, and no standard treatments with reliable survival and trans-differentiation of iPS cell lines into neurons was possible. This variability likely reflects clonal differences in iPS cell lines that are not immediately apparent in standard assays.

Although lentiviral Ngn2-induced trans-differentiation of iPS and ES cells into neurons is reproducible and reliable in an individual lab setting, it is not well suited for up-scaling. Therefore, FCDI optimized a large-scale manufacturing strategy to produce Ngn2-induced human neurons via the PiggyBac transposon system (see Methods). This system utilizes stable transgene integration by PiggyBac, which contains an inducible Ngn2 and a constitutive puromycin expression cassette. Puromycin-resistant cells were selected and expanded prior to doxycycline induction. Next, cryopreservation of Ngn2-induced human neurons was optimized, such that post-mitotic neurons can be cryopreserved in a fashion where they retain their pre-cryopreservation phenotype after thawing, and batches of cryopreserved human neurons were benchmarked against fresh cultures (see Methods). Lastly, post-thaw culture conditions were optimized so that Ngn2-induced human neurons can be cultured long-term in any laboratory CO_2_ incubator under defined media conditions and yield electrically and synaptically functional human neurons for imaging and whole-cell patch-clamp electrophysiology experiments (see Methods).

### Heterozygous *NRXN1* deletions (*NRXN1^del^*) predisposing to schizophrenia do not affect neuronal morphology or synapse numbers

We previously characterized the effect of conditional heterozygous *NRXN1* mutations on the molecular, cellular and electrophysiological properties of human neurons (Pak et al., 2015). In these studies, *NRXN1*-mutant neurons were derived from engineered ES cells carrying conditional heterozygous *NRXN1* deletions or truncations that could be produced by recombinases. We found that heterozygous *NRXN1* mutations did not impair dendritic arborization or synapse formation of human neurons, nor did they alter their passive membrane properties or action potential generation properties (Pak et al., 2015). Therefore, we tested the same parameters in Ngn2-induced neurons generated from pairs of schizophrenia patient-derived *NRXN1^del^* and control iPS cells. Neurons were either generated with the small-scale method and analyzed at Stanford, or were generated by the PiggyBac-based large scale method at FCDI and analyzed at Rutgers (Fig. 1A, S3).

In examining the first two pairs of *NRXN1*^del^ and control neurons, we did not detect any significant changes in neurite outgrowth, number of primary dendritic processes or dendritic branch points, or soma size (Fig. 2A-B, 2E-F). Moreover, the density and size of synapses were not altered in *NRXN1*^del^ neurons (Fig. 2C-D, 2E-F). Interestingly, despite different Ngn2-dependent neuronal induction strategies (lentiviral transduction vs. piggyBac; laboratory vs. industrial), different laboratories (Stanford vs. Rutgers) and independent analytical strategies (IntelliCount semi-automated quantification vs. Metamorph counting), we consistently observed no deficit in synapse formation in *NRXN1*^del^ neurons (Fig. 2E-F, Pak et al., 2015; Fantuzzo et al., 2017; see Methods). Thus, similar to previously examined engineered *NRXN1* mutations, patient-derived neurons with *NRXN1*^del^ CNVs do not display major morphological changes.

**Figure 2:**
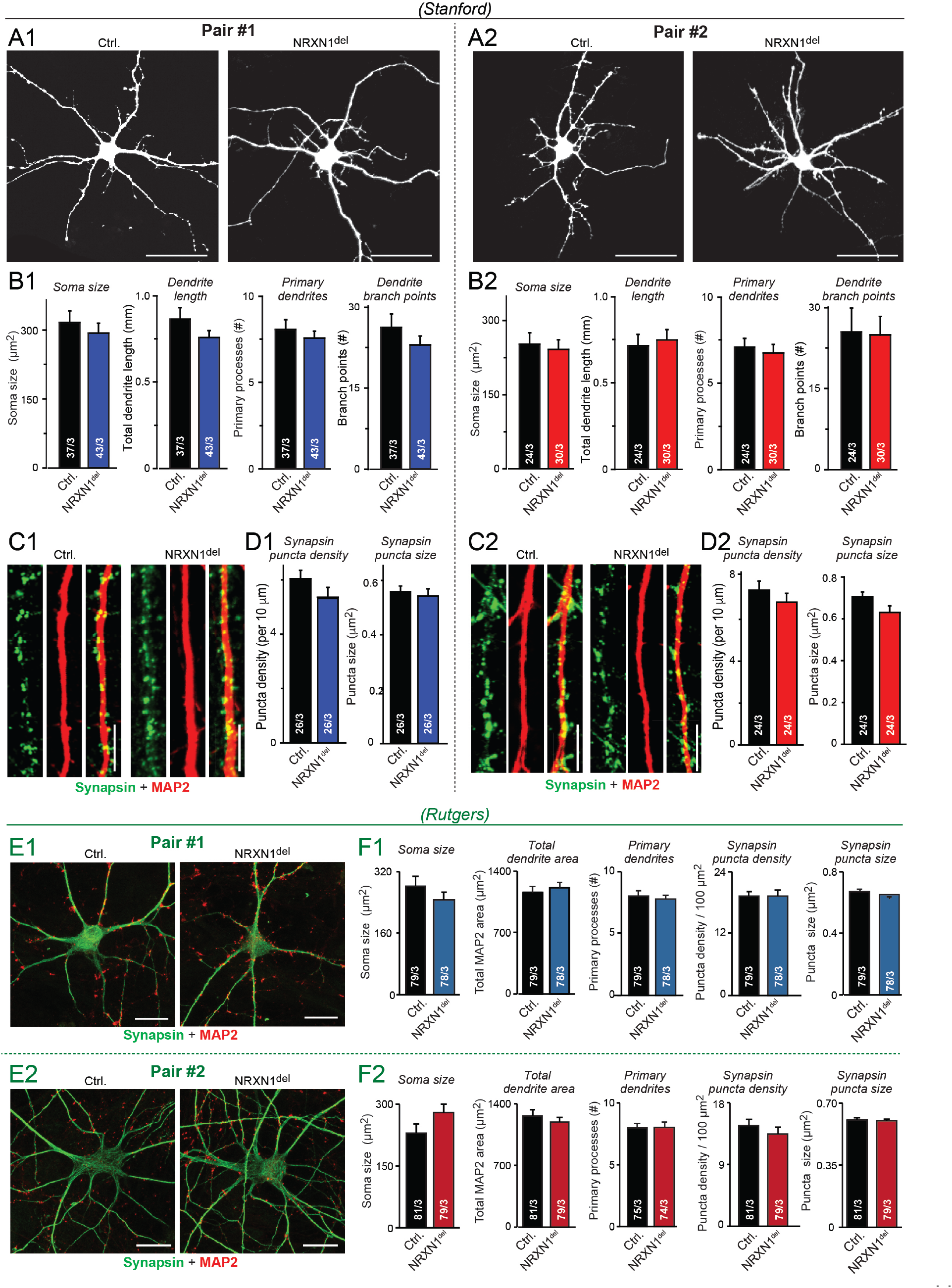
Dendritic outgrowth and synapse formation are not significantly altered in NRXN1^del^ neurons trans-differentiated from iPS cells derived from schizophrenia patients. **A** & **B**. *NRXN1^del^* neurons exhibit no changes in soma size or dendritic arborization as analyzed at Stanford University in neurons obtained from two pairs of patient-derived vs. control iPS cells (A1 & A2, representative confocal images of sparsely transfected cells expressing EGFP to label individual neurons for morphometric analyses [A1, A2, scale bar = 50 μm]; B1 & B2, summary graphs of soma size, dendrite length, number of primary dendrites, and dendritic branch points). Neurons were analyzed at 4 weeks in culture. **C** & **D**. *NRXN1^del^* neurons display no apparent change in synapse formation, assessed by the presynaptic marker Synapsin-1 as analyzed at Stanford University in two pairs of patient-derived vs. control neurons (C1 & C2, representative confocal images of MAP2-positive dendritic segments and Synapsin-1-positive synapses [scale bar = 5 μm]; D1 & D2, summary graphs of the density and size of Synapsin-1-positive puncta measured on secondary dendrites in control and schizophrenia-associated *NRXN1*^del^ neurons). **E** & **F.** Cross-lab and cross-platform validation confirming that *NRXN1^del^* neurons exhibit no significant changes in neuronal morphology or synapse numbers by analyses at Rutger’s University of the same two pairs of *NRXN1^del^* patient-derived vs. control neurons using different independent approaches. Specifically, neurons stained for MAP2 and Synapsin-1 were examined using the *Intellicount* automated quantification method (Fantuzzo et al., 2017) (E1 & E2, representative images [scale bar = 10 μm]; F1 & F2, summary graphs of the soma size, total dendritic tree area, number of primary processes, synapse density, and synapse size). Data are means ± SEM (numbers in bars represent number of cells/experiments analyzed). Statistical analyses by Student’s t-test comparing *NRXN1^del^* neurons to controls revealed no significant differences.

### Schizophrenia-associated *NRXN1^del^* CNVs do not significantly alter the intrinsic membrane properties of neurons

Next, we investigated the electrical properties of schizophrenia patient-derived *NRXN1*^del^ neurons. Using patch-clamp recordings, we measured the membrane capacitance, input resistance, neuronal excitability, resting membrane potential, and various parameters of action potential generation in two pairs of *NRXN1*^del^ and control neurons. We detected no differences between schizophrenia patient-derived *NRXN1^del^* neurons and corresponding control neurons in both pairs (Fig. 3), suggesting that heterozygous loss of *NRXN1* does not affect non-synaptic electrical properties of human neurons. Again, the data were consistent across the two different sites with neurons generated by different Ngn2 induction methods, providing a cross-platform validation of the results (Fig. 3).

**Figure 3:**
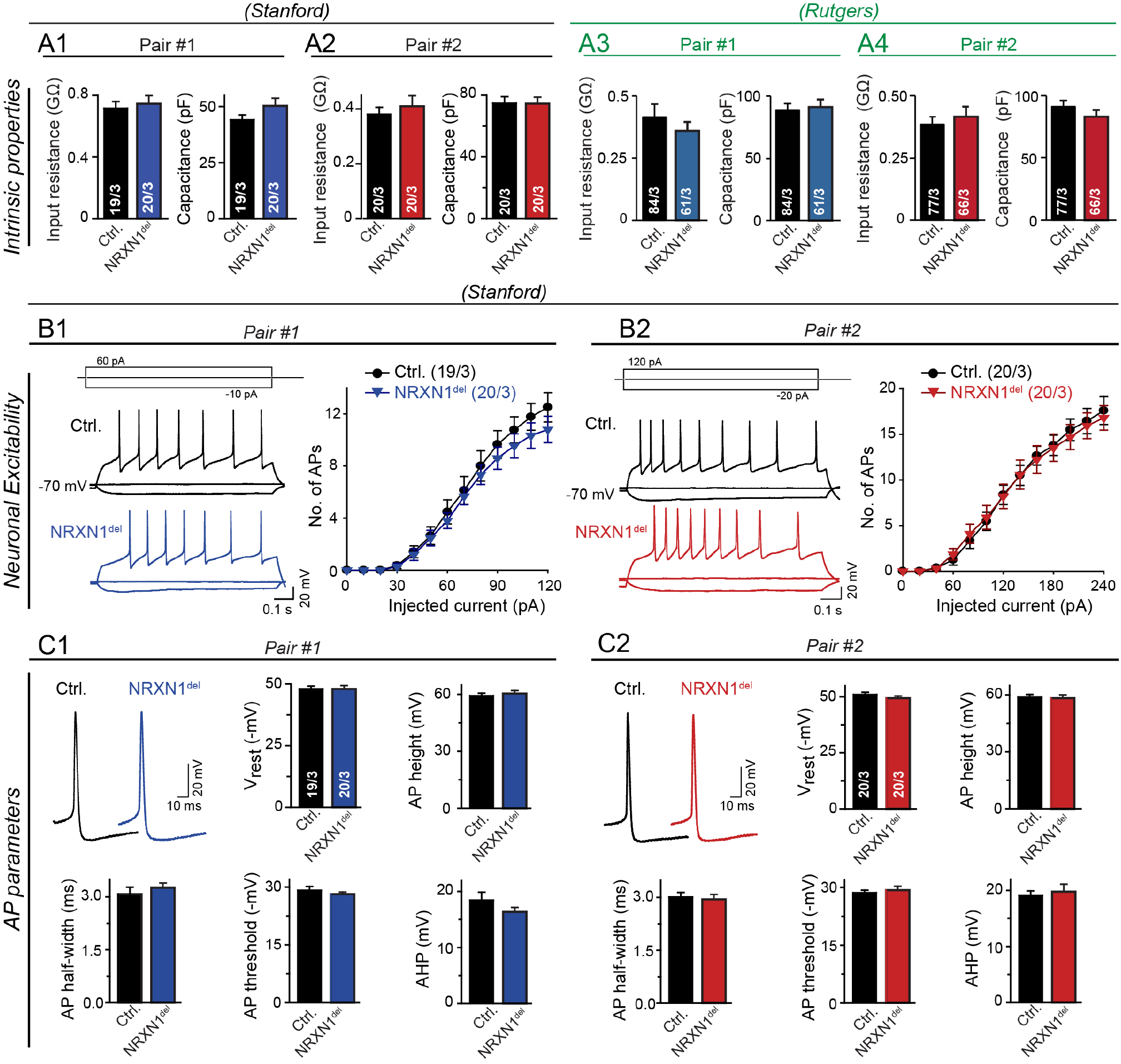
NRXN1^del^ neurons generated from schizophrenia patient-derived iPS cells exhibit no significant changes in intrinsic electrical properties. **A.** *NRXN1*^del^ neurons display no changes in passive membrane properties (left and right, summary graphs of the input resistance and capacitance, respectively) of Pair #1 (A1, A3) and Pair #2 (A2, A4) control and schizophrenia *NRXN1*^del^ neurons analyzed at Stanford University (A1, A2) and Rutgers University (A3, A4). **B.** *NRXN1*^del^ neurons display no changes in neuronal excitability as analyzed at Stanford University in Pair #1 (B1) and Pair #2 (B2) neurons (left, representative traces of action potentials induced by step current injections, illustration of experimental protocol is shown on top; right, intensity-frequency plots showing the number of evoked action potentials as a function of step current injections). **C.** *NRXN1*^del^ neurons exhibit no significant changes in action potential properties as analyzed in Pair #1 (C1) and from Pair #2 neurons (C2) at Stanford University (top left panels, representative current-clamp recording traces of individual APs; top right panels, summary graphs of the resting membrane potential [Vrest] and action potential height [AP height]; bottom panels, summary graphs of the action potential half-width [AP half-width], action-potential firing threshold [AP threshold], and action potential after-hyperpolarization potential amplitude [AHP]). Data are means ± SEM (numbers in bars represent number of cells/experiments analyzed). Statistical analyses by Student’s t-test comparing *NRXN1^del^* neurons to controls revealed no significant differences. All recordings were performed from 6 weeks after the start of differentiation.

### *NRXN1^del^* CNVs decrease the frequency of spontaneous synaptic ‘mini’ events

To analyze synaptic function, we asked whether *NRXN1*^del^ neurons exhibit a change in spontaneous miniature excitatory postsynaptic currents (mEPSCs, monitored in the presence of tetrodotoxin) or spontaneous EPSCs. Schizophrenia patient-derived *NRXN1^del^* neurons exhibited a large (~2-fold) decrease in mEPSC frequency, but no change in the mEPSC amplitude (Fig. 4). This decrease was observed in two independent pairs of *NRXN1*^del^ vs. control neurons, and was similarly detected in small-scale neurons produced and analyzed at Stanford, and in large-scale neurons produced at FCDI and analyzed at Rutgers. These results again phenocopy those previously obtained for engineered *NRXN1*-mutant human neurons (Pak et al., 2015), suggesting that the heterozygous *NRXN1* mutation impairs synaptic transmission.

**Figure 4:**
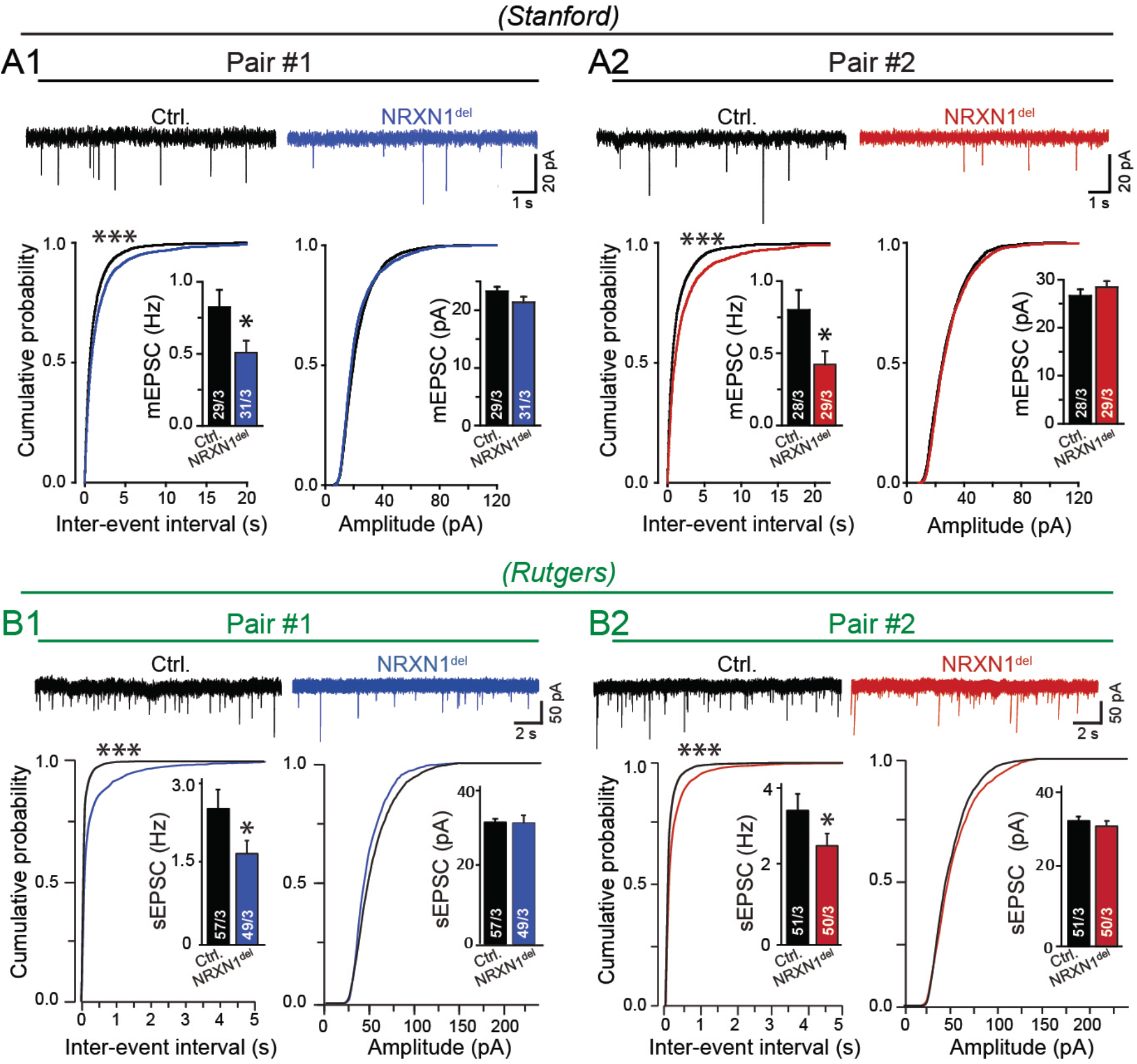
NRXN1^del^ neurons trans-differentiated from schizophrenia patient-derived iPS cells exhibit a significant decrease in the frequency of spontaneous excitatory synaptic events. **A.** The frequency but not amplitude of spontaneous mEPSCs is decreased ~2-fold in *NRXN1^del^* neurons derived from schizophrenia patients compared to controls (A1 and A2) (top, representative traces of mEPSC recordings; bottom left, cumulative probability plots of interevent intervals and summary graphs of the mEPSC frequency; bottom right, cumulative probability plots and summary graphs of the mEPSC amplitude). mEPSCs were recorded in the presence of tetrodotoxin (TTX, 1 μM) at Stanford University. **B.** Similar as A, but spontaneous EPSCs (sEPSCs) were analyzed in the absence of TTX with an independently generated set of neurons at Rutgers University (cross-lab and cross-platform validation). Data are means ± SEM (numbers in bars represent number of cells/experiments analyzed). Statistical analyses were performed by Student’s t-test for the bar graphs, and by Kolmogorov-Smirnov tests for cumulative probability plots, comparing *NRXN1^del^* to control neurons (* = p<0.05; non-significant comparisons are not indicated). All recordings were performed from 6 weeks after the start of differentiation.

### Schizophrenia-associated *NRXN1^del^* CNVs decrease the neurotransmitter release probability

To determine whether the decrease in mEPSC frequency reflects a decrease in synaptic strength and to determine whether this decrease may be caused by a change in neurotransmitter release probability, we measured action potential-evoked excitatory postsynaptic currents (EPSCs) mediated by α-amino-3-hydroxy-5-methyl-4-isoxazolepropionic acid receptors (AMPARs). Consistent with the decrease in mEPSC frequency, we observed a large decrease (~2-fold) in EPSC amplitude in *NRXN1^del^* neurons compared to controls both for the two pairs studied in the analyses described above, and also for a third pair of *NRXN1^del^* vs. control neurons (Fig. 5A).

**Figure 5:**
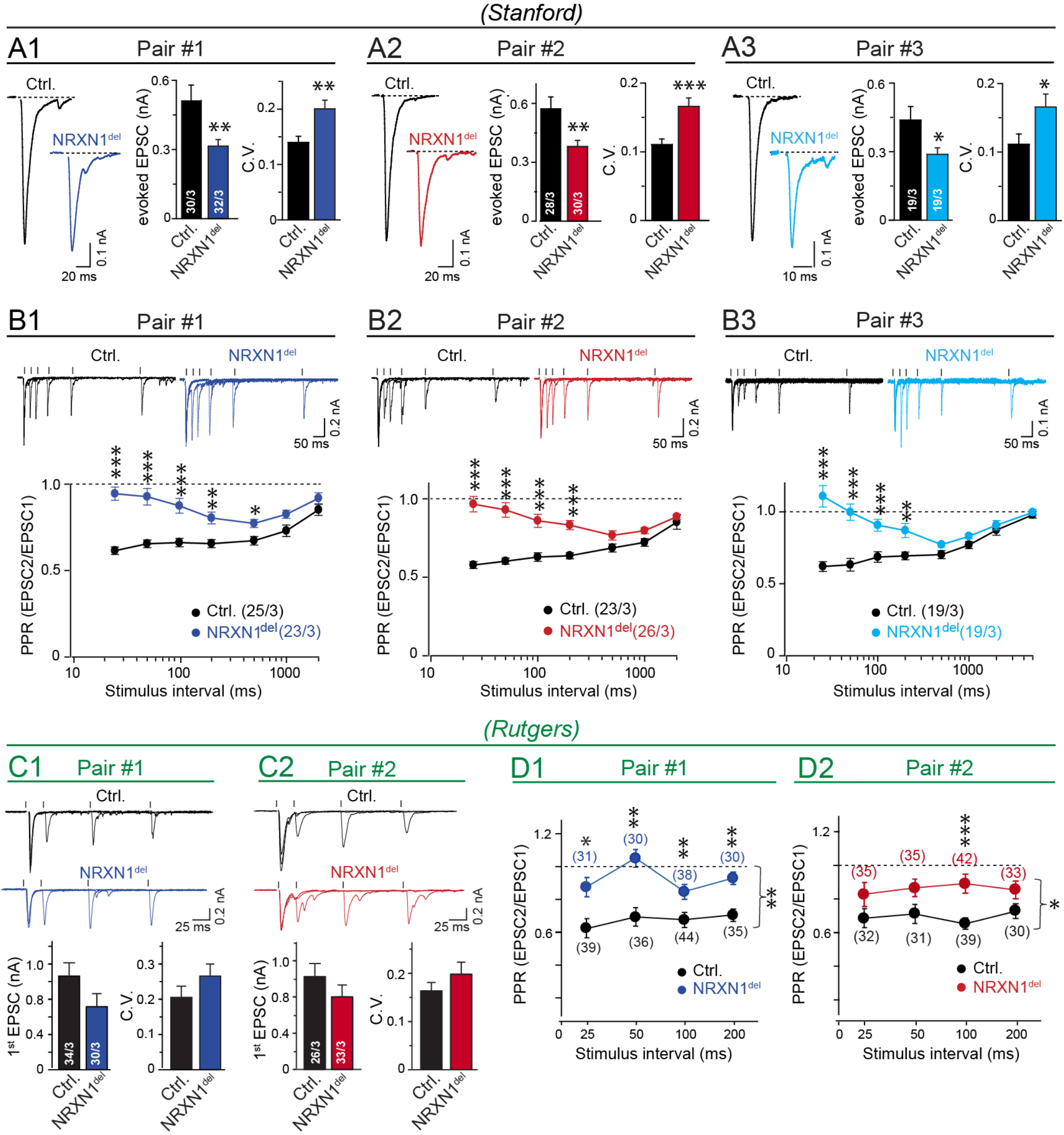
Evoked excitatory synaptic transmission is impaired in NRXN1^del^ neurons trans-differentiated from patient-derived iPS cells due to a decrease in release probability. **A.** *NRXN1*^del^ neurons display an ~30-40% decrease in the amplitude of evoked AMPAR-mediated EPSCs along with an increase in the coefficient of variation (C.V.) of evoked EPSC amplitudes in Pair #1 (A1), Pair #2 (A2) and Pair #3 (A3) of control and schizophrenia *NRXN1*^del^ neurons, as analyzed at Stanford University (A1-A3 = separate analyses of the three pairs of patient-derived vs. control neurons; left, representative traces of evoked EPSCs; middle and right, summary graphs of the EPSC amplitude and coefficient of variation [C.V.]). **B.***NRXN1*^del^ neurons display a significant increase in the paired-pulse ratio (PPR) of evoked AMPAR–mediated EPSCs compared to matched control neurons, as analyzed at Stanford University (B1-B3: separate analyses of the three pairs of patient-derived vs. control neurons; top, representative traces of PPR recordings, only the first 5 interstimulus interval is shown (25-500ms) and stimulus artifacts are removed for better overview; bottom, summary plots of PPR as a function of the interstimulus intervals in the range of 25-5000 ms). **C**& **D**. *NRXN1^del^* neurons derived from schizophrenia patients exhibit a decrease in the amplitude of evoked EPSCs and an increase in the PPR of EPSCs when compared to matched controls, as analyzed at Rutgers University for two patient-control pairs (C top, representative traces of EPSCs recorded in response to two sequential action potentials elicited with 25-200 ms intervals; C bottom, summary graphs of the amplitude and coefficient of variation [C.V.] of the first EPSC response evoked by the first of the two sequential action potentials; D, summary plot of the PPR of EPSCs elicited with 25-200 ms intervals). Data are means ± SEM (numbers in bars represent number of cells/experiments analyzed). Statistical analyses were performed by Student’s t-test for the bar graphs (A, C) and by two-way RM ANOVA with Tukey’s post-hoc test for summary plots (B, D), comparing *NRXN1^del^* neurons to controls (* = p<0.05; ** = p<0.01; *** = p<0.001; non-significant comparisons are not indicated). Neurons were analyzed at 6 weeks in culture.

We then examined whether the decreased synaptic strength in *NRXN1^del^* neurons is due to a decrease in release probability, since the lack of a change in mEPSC amplitudes suggested that postsynaptic AMPARs were normal. We assessed the release probability by measuring the coefficient of variation (C.V.) of evoked EPSCs, which is inversely proportional to the release probability (Hefft et al., 2002). The coefficient of variation of EPSCs was increased significantly (~1.4-fold) in *NRXN1^del^* vs. control neurons, suggesting a decrease in release probability (Fig. 5A). To independently test this conclusion, we measured the paired-pulse ratio (PPR) of EPSCs, which is the ratio of the EPSC amplitudes evoked by two closely spaced action potentials. The PPR depends on the release probability because the extent of release induced by the first action potential determines, among others, how much additional release can be induced by the second action potential (Abbott and Regehr, 2004; Neher and Brose, 2018). In control neurons, the PPR exhibited a decrease in the second response (referred to as paired pulse depression) because the release probability under our recording conditions is high (Fig. 5B). However, in the *NRXN1^del^* neurons generated from patient-derived iPS cells, the PPR was greatly increased (i.e., paired-pulse depression was decreased), confirming a deficit in the initial release probability. This increase in PPR was replicated in all three pairs of *NRXN1^del^* vs. control neurons (Fig. 5B).

We performed similar experiments with the two pairs of *NRXN1^del^* and control neurons that we primarily analyzed at Rutgers U., with comparable results (Fig. 5C, 5D). Again, we observed a decrease in the EPSC amplitude and an increase in the coefficient of variation of the EPSC, although the difference was not significant (Fig. 5C). Moreover, we detected an increase in the PPR, which was highly significant (Fig. 5D). Taken together, these results strongly indicate that patient-derived *NRXN1^del^* neurons exhibit a robust and reproducible decrease in neurotransmitter release probability.

### A newly engineered conditional *NRXN1^del^* iPS cell line reproduces the synaptic impairments observed in patient-derived *NRXN1^del^* neurons

In analyzing patient-derived neurons, a pressing question is whether the observed phenotypes in neurons with disease-associated CNVs or mutations are truly caused by these genetic changes, or are produced by polygenic effects resulting from a combination of a specific mutations with a particular genetic background (Hyman, 2015). Mutations associated with neuropsychiatric disorders often predispose to multiple disease conditions. For example, *NRXN1* CNVs are among the most frequent mutations associated with not only schizophrenia, but also intellectual disability, autism-spectrum disorders, epilepsy and others (Castronovo et al., 2020; Lowther et al., 2017, Kasem et al., 2018; Szatmari et al., 2007; Guilmatre et al., 2009; Liu et al., 2012; Ching et al., 2010; Nag et al., 2013; Huang et al., 2017). We thus decided to test whether the similarity of the synaptic impairments we observed in *NRXN1^del^* neurons generated from schizophrenia patient-derived iPS cells to those we previously described for *NRXN1*-deficient neurons generated from ES cells can be validated with neurons generated from iPS cells carrying a newly engineered conditional *NRXN1* mutation. For this purpose, we generated a new iPS cell line (from our control line C3141a, Table 1) carrying a heterozygous conditional KO (cKO) of *NRXN1* using a strategy that is identical to the approach we previously employed in mice and in human ES cells (Fig. S4, Table 2). We produced this additional validation tool not only to confirm our conclusions, but also to generate a conditionally mutant *NRXN1* iPS cell line that can be freely distributed for further studies, and is not subject to the commercial constraints imposed on the previously generated conditionally mutant *NRXN1* ES cell lines (Pak et al., 2015).

**Table 2.**
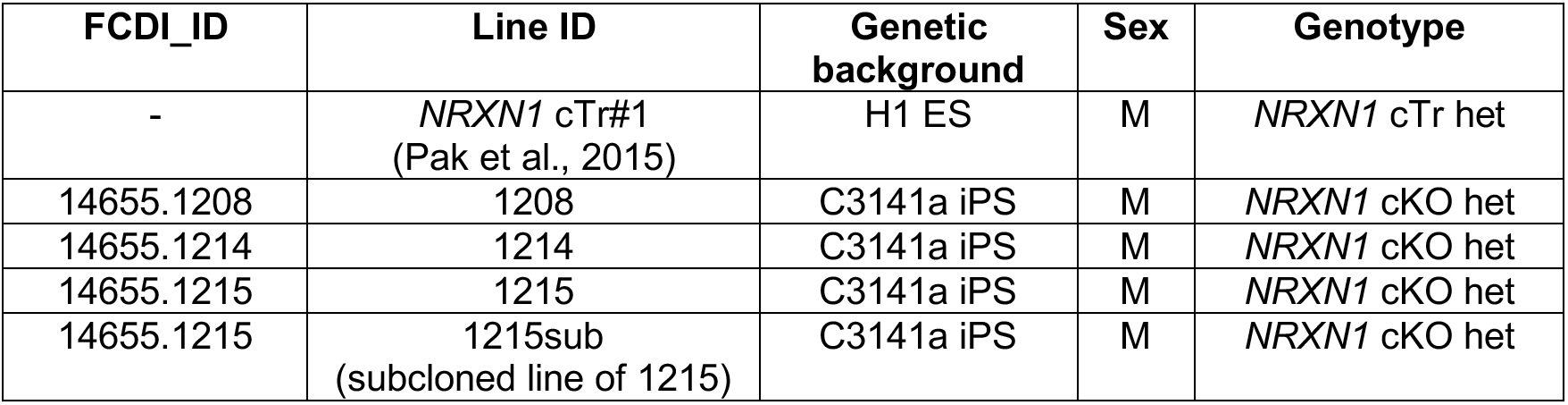
Engineered iPSC/hESC lines.

We limited our analysis of the newly engineered conditional *NRXN1^del^* neurons to key measurements, namely assessments of mEPSCs and evoked EPSCs. The newly engineered *NRXN1^del^* neurons exhibited the same decrease in mEPSC frequency and EPSC amplitude and the same increase in PPR as the patient-derived *NRXN1^del^* neurons (Fig. 6, S5A). These results confirm that heterozygous *NRXN1* deletions produce a robust synaptic phenotype in human neurons, suggesting that the associations of heterozygous *NRXN1* deletions with specific neuropsychiatric disorders must be shaped by factors downstream of the synaptic impairments. Since this newly engineered line was generated in an iPS cell line that can be distributed without commercial restrictions, this line is now widely available for further mechanistic and therapeutic studies in schizophrenia.

**Figure 6:**
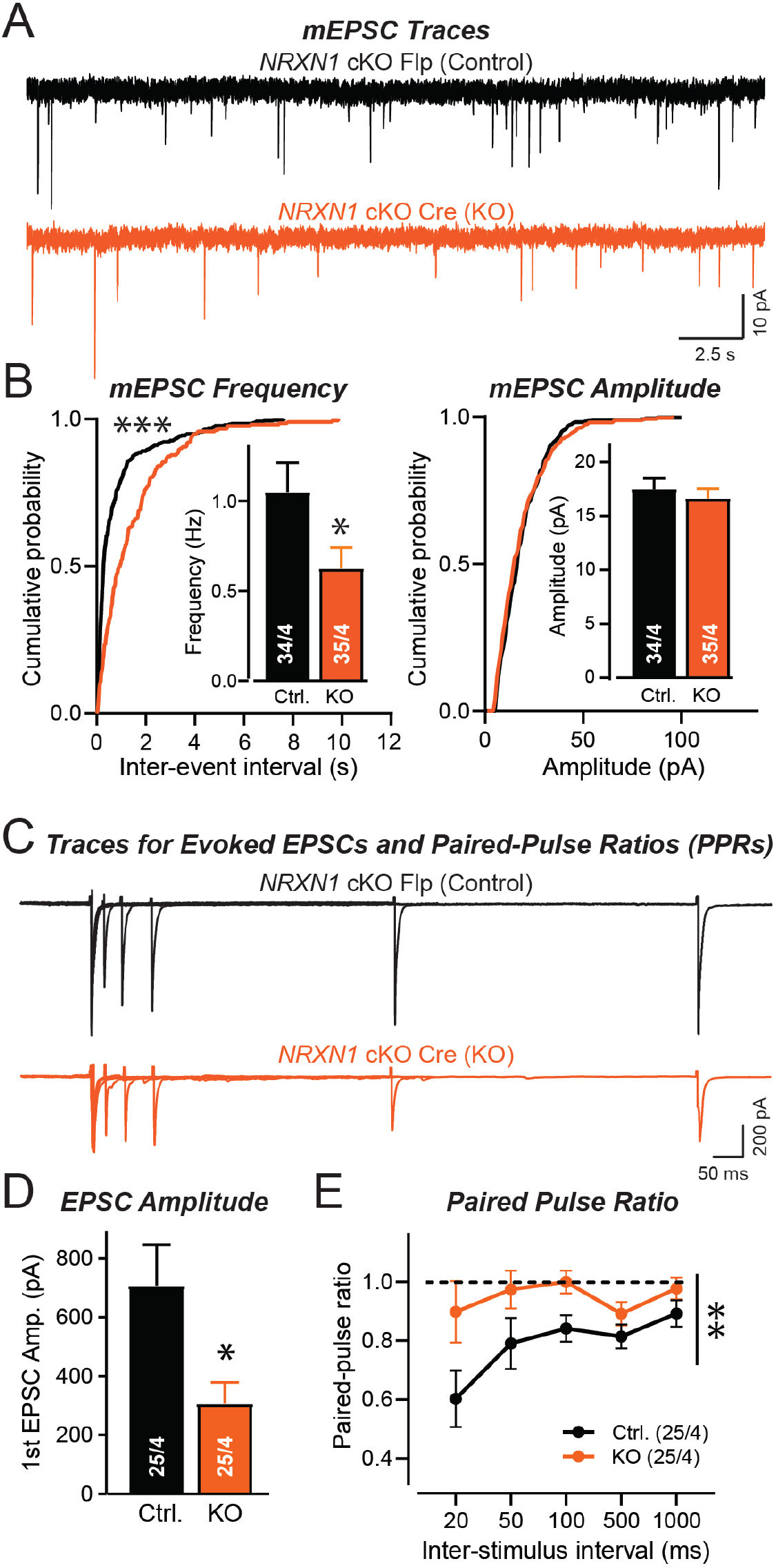
Heterozygous deletion of human NRXN1 using a newly engineered conditionally mutant iPS cell line replicates the impairment in presynaptic release probability observed in NRXN1^del^ neurons trans-differentiated from schizophrenia patient-derived iPS cells. **A.** Sample traces of mEPSCs recorded from human neurons derived from the newly engineered *NRXN1* cKO iPS cell line (1215sub). Neurons were infected with lentiviruses expressing Flp (creates wild-type genotype and serves as a control [black]) or Cre (induces heterozygous *NRXN1* KO [orange]). See Table 2 and Fig. S4 for a description of the newly engineered conditional *NRXN1* mutation. **B.** Conditional heterozygous *NRXN1* deletion in neurons generated from a newly engineered iPS cell line causes a nearly 2-fold decrease in mEPSC frequency but not amplitude (left, cumulative probability plot of the mEPSC interevent intervals and summary graph of the mEPSC frequency; right, cumulative probability plot and summary graph of the mEPSC amplitude). **C.** Sample traces of evoked EPSCs elicited by closely spaced pairs of action potentials (intervals: 20 ms, 50 ms, 100 ms, 500 ms, 1000 ms) to measure both the EPSC amplitude and the EPSC paired-pulse ratio from neurons derived from the *NRXN1* cKO iPS cell line. **D.** Conditional heterozygous *NRXN1* deletion in engineered neurons induces a >2-fold decrease in the amplitude of evoked EPSCs (summary graphs of the amplitude of the first evoked EPSC). **E.** Conditional heterozygous *NRXN1* deletion in engineered neurons causes a >2-fold reduction in the paired-pulse depression of evoked EPSCs (summary plot of the paired-pulse ratio (PPR), measured as the amplitude ratio of the second over the first evoked EPSC, and plotted as a function of the inter-stimulus interval). Data are means ± SEM; numbers of cells/cultures analyzed are shown in the bar diagrams. Statistical significance was evaluated with the Kolmogorov-Smirnov-test (cumulative probability plots), one-tailed t-test (summary graphs), or two-way ANOVA (PPR plots) (*, p<0.05; **, p<0.01). All recordings were performed from 6 week-old neurons.

### Mouse neurons carrying the heterozygous *NRXN1^del^* allele do not recapitulate the human *NRXN1^del^* phenotype

In our earlier analysis of the engineered conditional human *NRXN1^del^* neurons, we found that dissociated cultures of mixed mouse neurons and glia obtained from the cortex of heterozygous or homozygous *Nrxn1α* KO mice did not replicate the heterozygous *NRXN1^del^* phenotype (Pak et al., 2015). This appeared to indicate that human neurons are uniquely sensitive to the heterozygous deletion of *NRXN1*, which is unexpected given the apparent structural and functional similarity between human *NRXN1* and mouse *Nrxn1*. However, in these original experiments the compared human and mouse neurons were quite different. The mouse neurons were primary cultures of many cell types that were analyzed as a mixture derived from the mouse cortex after neurogenesis and neuronal migration had completed, whereas the human neurons constituted a relatively homogeneous population of directly trans-differentiated neurons. Moreover, the mouse neurons were from *Nrxn1α* KO animals, whereas the human neurons carried heterozygous deletions of both *NRXN1α* and *NRXN1β*. Thus, these earlier experiments did not reveal whether or not heterozygous *Nrxn1* deletions in mouse neurons would produce a phenotype similar to that observed for the human neurons analyzed here.

To directly test whether human and mouse neurons are indeed differentially sensitive to heterozygous *NRXN1/Nrxn1* deletions, we repeated the precise experiment that we had performed with engineered human *NRXN1^del^* neurons with the equivalent mouse *Nrxn1^del^* neurons. For this purpose, we generated ES cells from mice with exactly the same mutation as the human conditional *NRXN1^del^* iPS and ES cells (Chen et al., 2017; Trotter et al., 2019). We converted these mouse ES cells into Ngn2-induced neurons, and analyzed the electrophysiological phenotype of these neurons. Strikingly, the mouse neurons did not exhibit a significant synaptic impairment (Fig. 7, S5B). They displayed a normal mEPSC frequency, and a trend towards a decrease in EPSC amplitude that was not statistically significant (Fig. 7A-7D). Moreover, the mouse *Nrxn1^del^* neurons showed no change in paired-pulse ratio, possibly the most sensitive electrophysiological phenotype (Fig. 7E). These results indicate that the synaptic phenotypes conferred by the heterozygous human *NRXN1^del^* mutations are unique to the human neuronal context.

**Figure 7:**
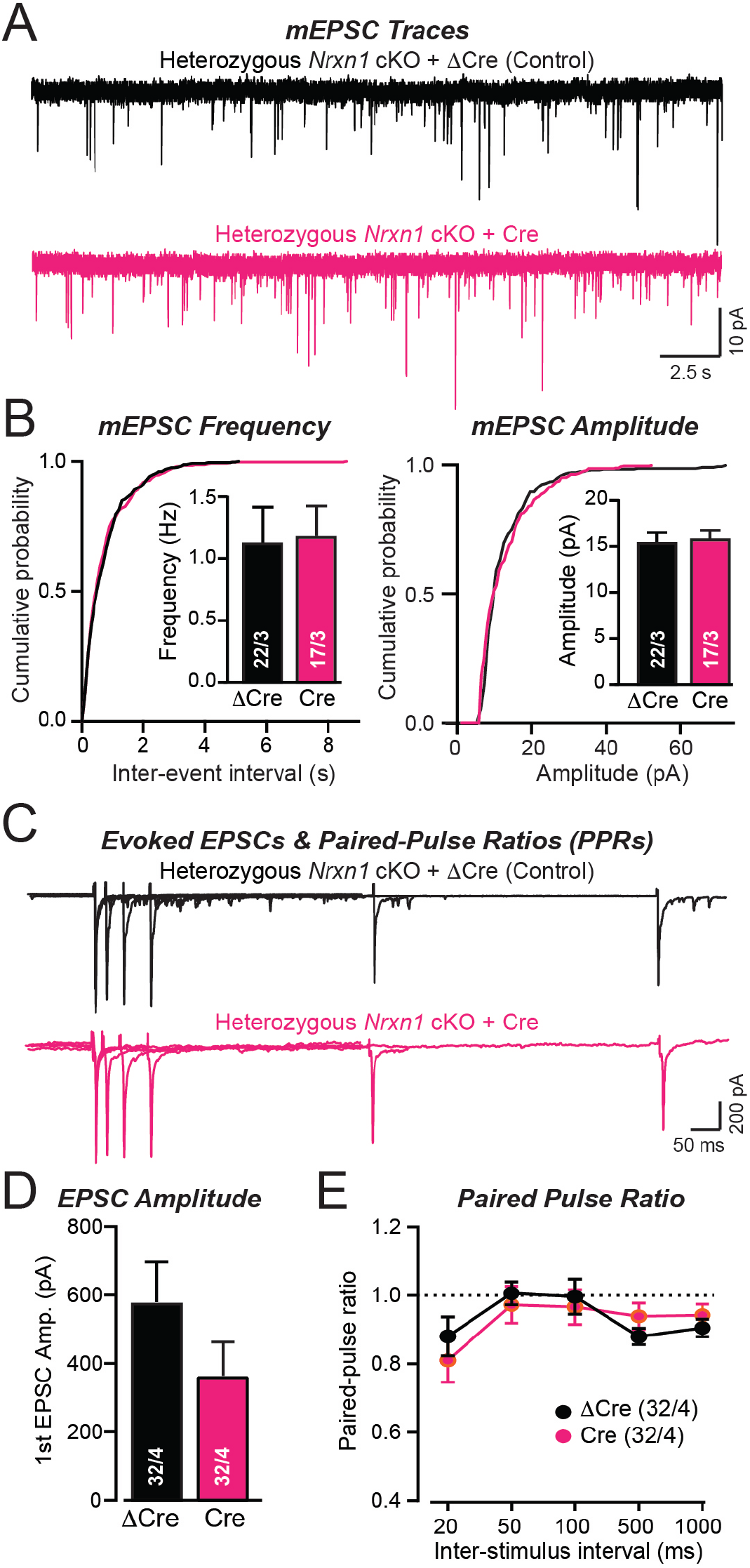
Mouse Nrxn1^del^ neurons fail to replicate the impairment in presynaptic release probability of human NRXN1^del^ neurons. **A** & **B**. Heterozygous *Nrxn1* deletions in mouse neurons cause no significant changes in mEPSC frequency or amplitude. ES cells isolated from heterozygous *Nrxn1* cKO mice (Chen et al., 2017) were trans-differentiated into neurons using Ngn2 expression, infected with lentiviruses expressing Cre (to induce the heterozygous *Nrxn1* deletion) or ΔCre (control), and analyzed at DIV14-16 (A, representative mEPSC traces; B, cumulative probability plots of the mEPSC interevent intervals [inset = summary graph of the mEPSC frequency] and of the mEPSC amplitude [inset = summary graph of the mEPSC amplitude]). **C**-**E**. Heterozygous *Nrxn1* deletions in mouse neurons also cause no significant impairments in evoked synaptic transmission. EPSCs were recorded in response to two sequential stimuli separated by defined inter-stimulus intervals (20 ms, 50 ms, 100 ms, 500 ms, 1000 ms) in neurons prepared as decribed in A (C, representative traces; D, summary graphs of the amplitudes; E, summary plot of the paired-pulse ratio of the EPSCs displayed as a function of the interstimulus intervals). Numerical data are means ± SEM (number in bars or brackets show number of cells/cultures analyzed). No significant differences were detected between test and control conditions using an unpaired, one-tailed t-test (B & D) or a two-way ANOVA (E).

### Heterozygous *NRXN1* deletions consistently increase CASK protein levels in human neurons

One striking finding previously obtained in ES cell-engineered *NRXN1^del^* neurons was the increase in CASK protein stability, which was detected with two different engineered *NRXN1* mutations (Pak et al., 2015). CASK is a cytoplasmic scaffolding protein that interacts with neurexins (Hata et al., 1996) and forms a tight complex with other presynaptic proteins (Wei et al., 2011; Cohen et al., 1998; Hsueh et al., 1998; Butz et al., 1998; Tabuchi et al., 2002). In human patients, mutations in *CASK* – which is X-linked– are associated with autism spectrum disorders and X-linked mental retardation in addition to brain malformations (Najm et al., 2008; Saitsu et al., 2012; Piluso et al., 2009; Hackett et al., 2010; Sanders et al., 2012; Neale et al., 2012). We thus tested whether CASK protein is also increased in schizophrenia patient-derived *NRXN1^del^* neurons. Consistent with previous studies, we observed a large increase (>50%) in CASK protein levels in *NRXN1^del^* neurons, while all other synaptic proteins measured were not altered, with the exception of *NRXN1* protein levels, which were downregulated by ~50% (Fig. 8A, 8B). Moreover, we detected a similar increase in CASK levels in neurons derived from the newly engineered heterozygous *NRXN1^del^* iPS cell line (Fig. 8C, 8D), further validating this change. Since no change in CASK mRNA levels was detected in the RNAseq experiments (see below), these results indicate that CASK protein is stabilized and protected from degradation upon heterozygous deletion of *NRXN1* in human neurons.

**Figure 8:**
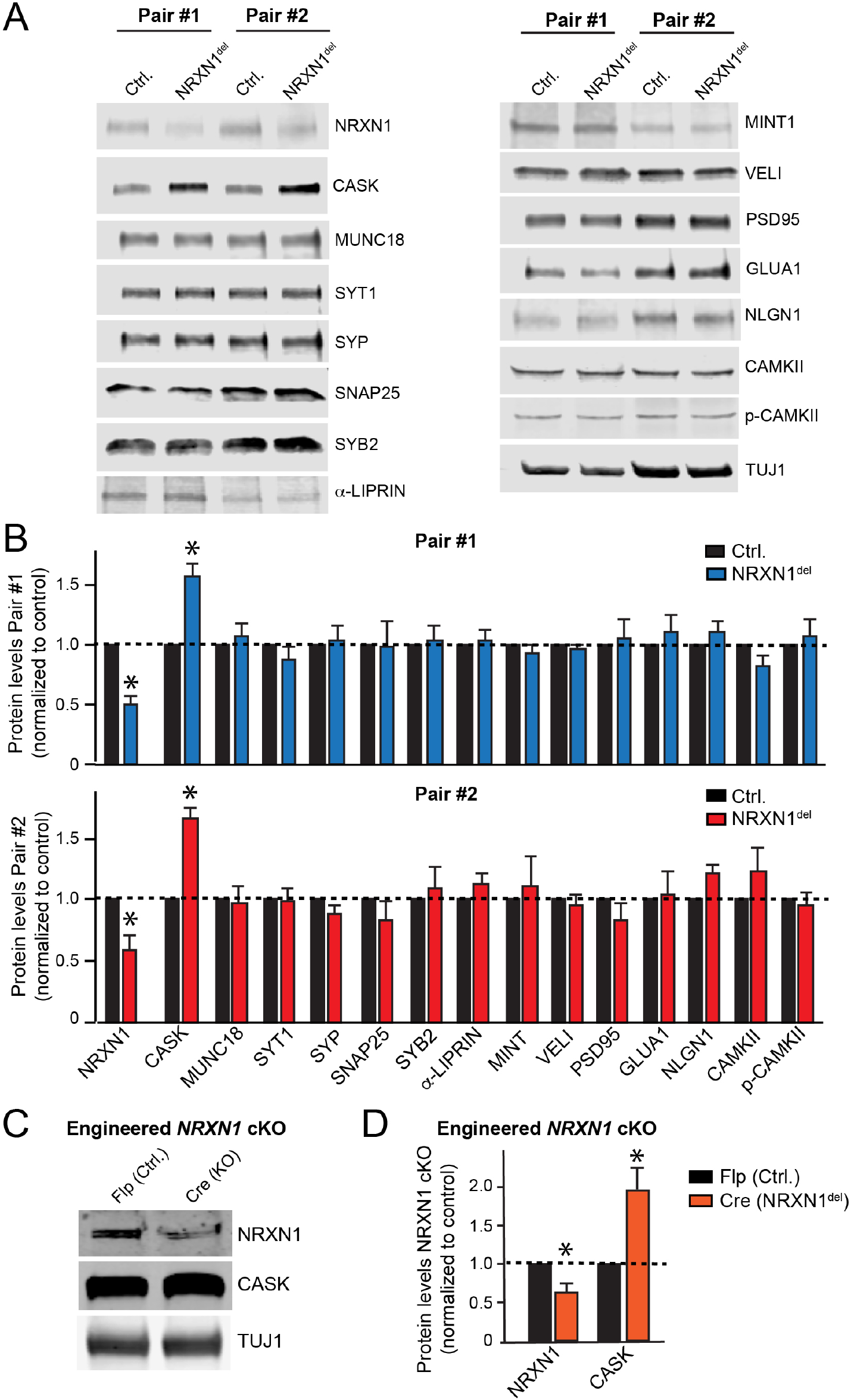
CASK protein levels are up-regulated in NRXN1^del^ neurons generated from schizophrenia patient-derived iPS cells and from genetically engineered iPS cells. **A.** Representative immunoblots of neurons generated from two pairs of patient-derived *NRXN1^del^* vs. control iPS cells. Neurons were analyzed at 4 weeks in culture (abbreviations: SYT1, Synaptotagmin-1; SYP, Synaptophysin; SYB2, Synaptobrevin-2; p-CAMKII, CaMKII-Phospho-286/7). **B.** Quantifications of protein levels in the two pairs of *NRXN1^del^* and control neurons reveal that apart from a large increase in CASK protein levels and a decrease in neurexin levels, no other major changes were detected in the levels of the analyzed proteins. All analyses were carried out using quantifications with fluorescently labeled secondary antibodies and TUJ1 as a loading control. **C**& **D**. Representative blots (C) and quantifications (D) reveal that the engineered conditional *NRXN1* deletion in an iPS cell background (1215sub) also causes an increase in CASK protein levels. Experiments were performed as in A & B. Data are means ± SEM (n = 3-5 independent cultures). Statistical analyses were performed by Student’s *t* test comparing test samples to the control (\**p*<0.05).

### *NRXN1^del^* neurons exhibit modest characteristic changes in gene expression

Using four-weeks old, relatively mature human neurons generated from the three pairs of patient-derived *NRXN1^del^* and control iPS cells analyzed in Fig. 5, we performed bulk RNA-sequencing (RNAseq) on total RNA in triplicate (Fig. 9A). In addition, we performed bulk RNAseq analyses (in triplicate) on isogenic pairs of human neurons without or with the heterozygous *NRXN1* deletion that were trans-differentiated from the newly engineered iPS cell line carrying a conditional *NRXN1^del^* allele (Fig. 6, S4), and on 3 unrelated wild-type iPS cell lines (in duplicate). In total, we performed differential gene expression analyses on 30 samples (9 controls vs. 9 schizophrenia-*NRXN1* mutants, 3 controls vs. 3 engineered *NRXN1* mutants, and 6 wild-type iPS cell samples) in triplicates or duplicates (replicates refer to independent cultures performed at different time points).

**Figure 9:**
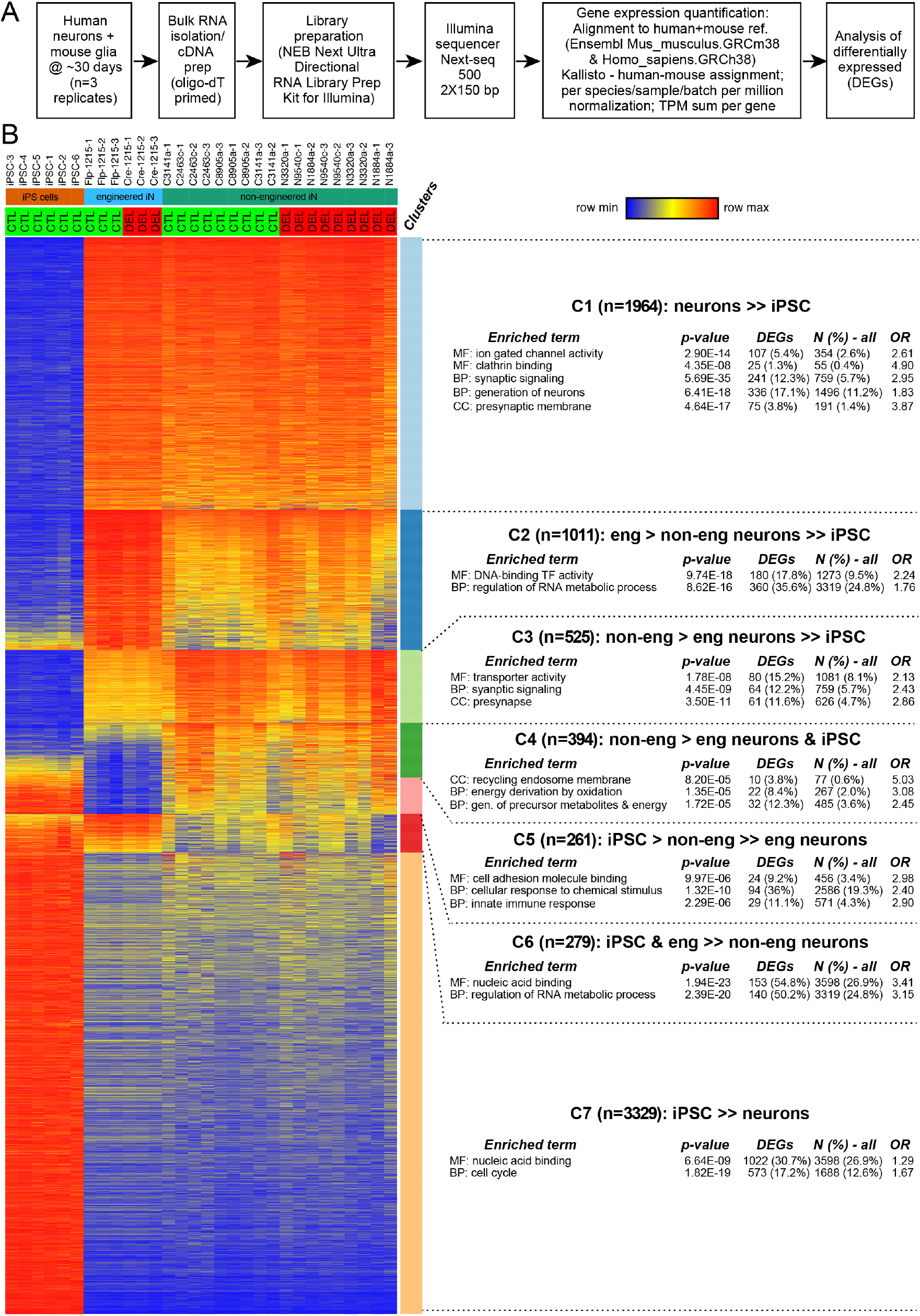
RNAseq analyses comparing the transcriptomes of control iPS cells and of NRXN1^del^ and control neurons generated from patient-derived, control, and engineered conditional iPS cells reveal distinct patterns of differential gene expression. **A.** Flow-diagram of RNAseq experiments. RNA-seq libraries from total RNA were prepared induced human neurons (~DIV28) co-cultured with mouse glia (triplicate cultures for all iN cell cultures; duplicate for iPS cell lines), using NEBNext Ultra Directional RNA Library Prep Kit for Illumina (NEB). Paired-end sequencing (150 base-pair reads) was performed using the Illumina Next-seq 500 platform (Novogene). RNA-seq reads were aligned to human + mouse reference (Ensembl Mus_musculus. GRCm38 and Homo_sapiens. GRCh38 transcriptomes v94) using the stranded library option of Kallisto v0.46.0 (Bray et al., 2016) and a concatenated mouse-human index. From each Kallisto-processed mouse-human RNAseq sample, an expression matrix was constructed to generate human-specific TPMs (transcripts per million) by (1) counting the per million human reads per mouse-human sample; (2) summing up transcript-level TPM values per gene; and (3) converting gene TPM values to log2(TPM+1) values. 19,701 genes were selected with log2(TPM+1) ≥ 1 in at least one of the thirty samples, and the resulting gene-level values were per-sample quantile-polished to reduce sample-sample variability (Zyprych-Walczak et al., 2015). Genes for subsequent analyses were selected from 17,004 gene IDs with log2(TPM+1) ≥ 2 in at least one of the thirty samples. **B.** Cluster analysis of differentially-expressed genes (DEGs). For 15,000 post-QC genes meeting expression criteria (log[tpm+1] ≥ 2 in at least one 1 culture; and > 0.1 in all cultures) in one or more types of cells (non-engineered neurons [18n], engineered neurons [6e], iPS cells [6p]), four *limma-trend* DEG analyses were carried out (18n, 6e or all 24 neuronal vs. 6 iPS cell cultures; 18n vs. 6e). 7763 DEGs were selected (|t| ≥ 15 in any of the first three analyses and/or |t| ≥ 4 in the fourth). K-means clustering assigned DEGs to 11 clusters, collapsed to 7 based on cluster similarity (containing 1964, 1011, 525, 394, 261, 279 and 3329 genes respectively). ToppFun (ToppGene.cchmc.org) then determined how many of the 15,000 genes and of the DEGs in each cluster were annotated to each Gene Ontology term (in all categories: MF [Molecular Function], BP [Biological Process] and CC [Cellular Component]). Terms containing at least 10 of 15,000 genes were retained, and the 13,392 genes mapped to at least one of those terms were considered the Test Set. For each term, a hypergeometric p-value was computed to determine the probability of the observed N (or more) DEGs being annotated to the term, given the proportion of all 13,392 Test Set genes assigned to that term (see Supplementary Data for detailed results). The ****heat map**** shows the relative expression level of each gene for each culture. The **annotation** lists results for selected terms with the most significant p-values for each Cluster: the p-value; the number and percentage of cluster DEGs and of all Test Set genes assigned to the term, and the odds ratio (OR). Because GO categories annotate the same genes multiple times, the 5% alpha level for Bonferroni correction considered the number of tested terms in each category separately (MF: 1209, p=4.14E-05; BP: 6405, p=7.81E-06; CC: 846; 5.91E-05).

As a result of the co-culturing scheme of human neurons with mouse glia, the RNAseq data on neurons were composed of a mixture of human neuronal and mouse glia transcriptomes. A key step in processing of the RNAseq data was to deconvolve the two transcriptomes and to normalize the relative abundance of each mRNA from each species to the total number of mRNAs from that species only. As described in the Methods, the Kallisto program was able to unambiguously assign each paired 150 basepair sequence to mouse and human reference transcriptomes (Bray et al., 2016). The mouse:human mRNA ratios differed between cultures of neurons trans-differentiated from various iPS cell lines. To control for these differences, we adjusted each sample’s species-specific mRNA abundance based on Transcripts per Million (TPM) for each gene, summed these for each gene, and then carried out per-sample quantile normalization steps for each sample (Zyprych-Walczak et al., 2015) (Fig. 9A). This approach provided a reproducible abundance measure for each gene in each sample. The resulting values were used to form a log2(TPM+1) gene-by-sample matrix that was used for differential gene expression analyses.

As described above, our samples included three types of reprogrammed cells: induced neurons generated from iPS cell lines from patient and control donors; isogenic neuronal cultures generated from an engineered iPS cell line carrying a conditional heterozygous *NRXN1^del^* knockout construct; and wild-type iPS cells. To gain a better understanding of the transcriptomic state of our cultures, we selected 7,763 differentially expressed genes (DEGs) that met at least one of the following four criteria in t-tests that compared cultures of different types of cells: an absolute value of t ≥ 15 in comparisons of engineered vs. iPS cells, of non-engineered vs. iPS cells, or of induced neurons in general vs. iPS cells; or an absolute value of t ≥ 4 for engineered vs. non-engineered neurons. These tests used the *limma-trend* procedure (see Methods). K-means cluster analysis of these DEGs revealed 11 clusters which were reduced to 7 clusters based on similarity. Analyses of functional enrichment of Gene Ontology (GO) categories (Fig. 9B; see Methods for details) suggested that the different clusters were associated with different biological categories, but could not be assigned to defined functional biological pathways. For example, clusters C1 and C3 are both associated with ‘synaptic signaling’ and ‘presynapse’, whereas clusters C6 and C7 are both associated with ‘nucleic acid binding’.

We quantitatively compared the transcriptomes among the various groups of samples. Principal component analysis identified the most significant gene expression differences between the transcriptomes of iPS cells vs. induced neurons (Fig. S6). This result is consistent with the profound changes in gene expression induced by trans-differentiation of iPS cells into neurons (Vierbuchen et al., 2010; Pang et al., 2011; Wapinsky et al., 2013; Zhang et al., 2013). The second most significant gene expression differences were detected between the transcriptomes of neurons derived from control and patient-derived *NRXN1^del^* iPS cells vs. neurons derived from engineered iPS cells (independent of whether they carried a heterozygous *NRXN1* deletion or were wild-type) (Fig. S6). Neurons from patient-derived and control iPS cells exhibited an overall more mature neuronal gene expression signature compared to neurons from engineered iPS cells (Flp vs. Cre). Owing to this difference, sub-comparisons (e.g., comparing wild-type neurons from control vs. engineered iPS cells) also uncovered significant differences in gene expression. The large transcriptome differences between neurons from engineered vs. non-engineered neurons is striking because the engineered iPS cells were derived from one of the three control iPS cell lines, and thus have the same genetic background. The engineered iPS cells were subjected to stringent quality control such as karyotyping, suggesting that the transcriptome differences are not due to large chromosomal arrangements (Fig. S2). The process of subcloning and engineering iPS cells may thus be sufficient to produce large shifts in iPS cell properties, resulting in fundamental gene expression changes between neurons derived from these iPS cells. On the background of these gene expression differences, the strong similarity in the synaptic phenotypes of patient-derived and engineered *NRXN1^del^* neurons appears even more compelling.

A primary goal of the bulk RNAseq experiments was to assess the specific effect of heterozygous *NRXN1* deletions on gene transcription. No differences between *NRXN1^del^* and control neurons could be detected in the principal component analysis (Fig. S6), and none of the clusters of DEGs was driven by gene expression differences between *NRXN1^del^* vs. wild-type genotypes (Fig. 9B, S7). These observations indicate that the heterozygous *NRXN1* deletion does not produce major perturbations of gene expression, but do not rule out more subtle effects. To achieve a more granular analyses, we selected (after initial QC, see Methods) 12,910 genes with expression values ≥2 in at least one of 18 patient and control samples, and >0.1 in all 18, and used a moderated paired t-test (with *limma-trend*), pairing the patient and control replicates that were cultured in the same plate at the same time, to identify DEGs with the highest significance.

First, we examined the most strikingly dysregulated genes (see Supplementary Data for detailed results). No p-values were significant after Bonferroni correction, and only 15 DEGs achieved a Benjamini & Hochberg False Discovery Rate of 30%. We constructed a volcano plot highlighting 78 genes with uncorrected p ≤ 0.001 and/or a |log2(fold-change)| ≥ 0.8) (Fig. 10A) to consider whether any gene expression changes were plausibly caused by the *NRXN1* deletion (Fig. 10A, S8). No obvious functional signature emerged from these DEGs. For example, although the heterozygous *NRXN1* deletion produced a selective and large impairment in synaptic transmission, only two genes among the top DEGs encode specifically synaptic proteins (CPLX2 and C1QL3), both of which are downregulated (Fig. 10A). Note that *CASK* expression is not significantly upregulated in *NRXN1^del^* neurons despite the increase in CASK protein levels, confirming previous data that *CASK* is upregulated post-transcriptionally in *NRXN1^del^* neurons (See Supplementary Data, Pak et al., 2015).

**Figure 10:**
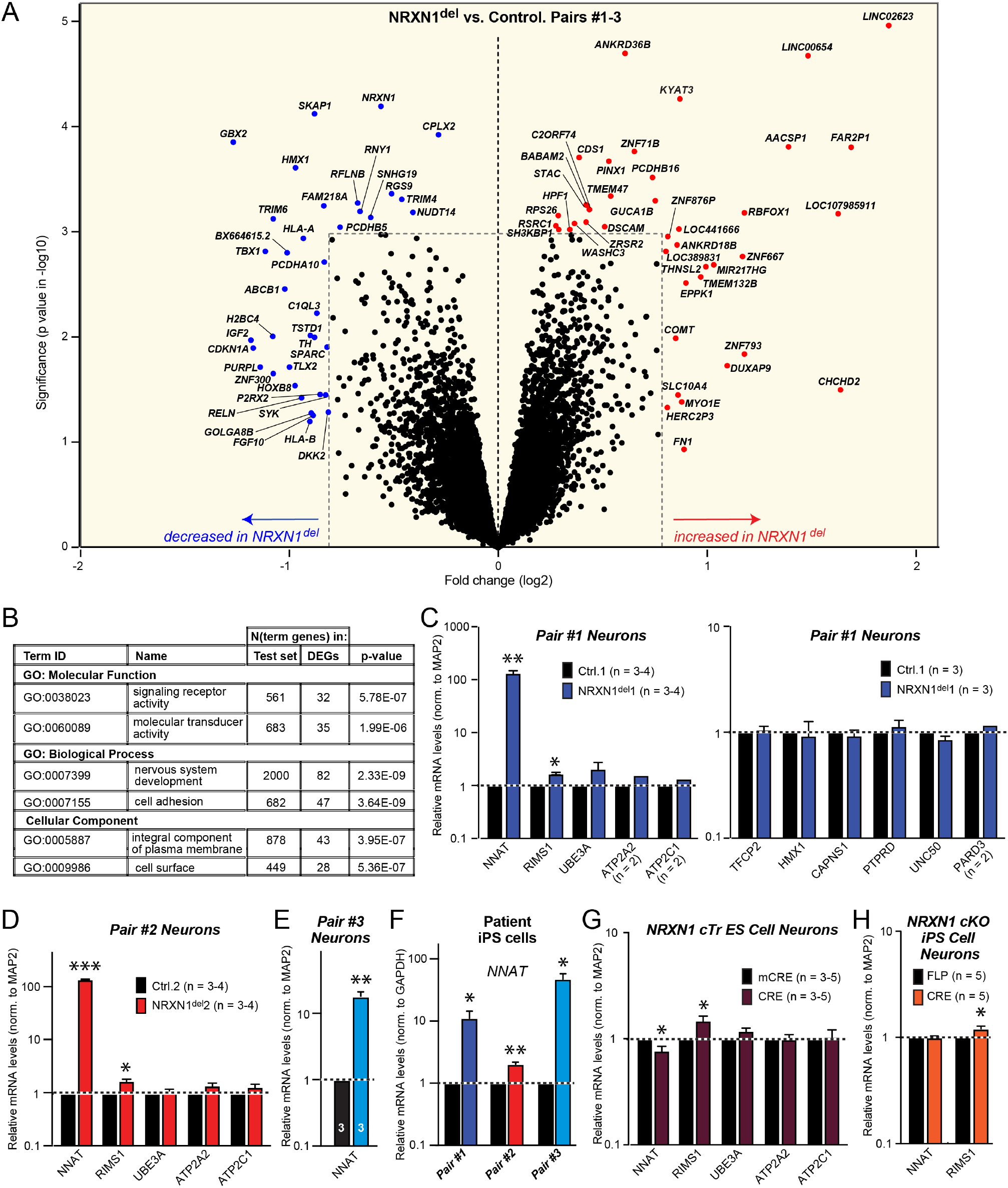
Gene expression is dysregulated in NRXN1^del^ neurons trans-differentiated from schizophrenia patient-derived iPS cells. **A**& **B.** For differential gene expression analysis of cultures of human neurons induced from patient-derived *NRXN1*^del^ and control iPS cells (N=9 in each group, Table 1), we selected 12,910 genes meeting all QC criteria (see Methods) including log2(TPM+1) expression values ≥ 2 in at least one of the 18 cultures and >0.1 in all cultures, and excluding small non-poly-adenylated RNA genes; alternative spliced, readthrough and fusion genes to minimize mapping errors. Moderated paired t-tests were performed using *limma-trend*, pairing the patient and control culture that were generated at the same time on the same plate. We examined these results in two complementary ways. **A.** The volcano plot shows p-value in negative logarithmic scale and log2(fold-change) results for all genes, highlighting those with the largest observed effects: |log2(fold-change)| ≥ 0.8 and/or -log10(P-value) ≥ 3. **B.** Shown are the two most significant Gene Ontology terms in each GO category, from functional enrichment analysis of the patient vs. control DEGs. A “broad” set of DEGs (403 up-regulated and 296 down-regulated) was selected with |log2(fold-change)| > 0.2 *and* p-value < 0.05, for this analysis (see Supplementary Data for a list of all DEGs) performed with ToppFun (ToppGene.org). There were 12,880 genes with a valid Gene Symbol, of which 11,781 (the Test Set) were annotated to at least one GO term, including 338 up-regulated and 257 down-regulated DEGs. Separately for up- and down-regulated DEGs, and for each term containing 10-5,000 Test Set genes, we performed a hypergeometric test (to determine the probability, given n DEGs and m Test Set genes in the term, of observing n or more of 257 or 338 DEGs with the expectation of m of 11,781 Test Set genes). For each main category, the 5% significance threshold was Bonferroni-corrected for the N of terms in that category (1,105 for Molecular Function; 5,962 for Biological Process; 807 for Cellular Component). There were no significant terms for upregulated DEGs. For down-regulated DEGs, there were 3, 32 and 10 significant terms respectively for the three categories; for a 5% Benjamini-Hochberg FDR threshold, there were 12,278 and 34 terms (note there is substantial overlap among terms, making the test conservative). **C**-**E**. *NNAT* expression is increased 10-100 fold in *NRXN1^del^* neurons derived from three schizophrenia patients. Relative expression levels of the indicated mRNAs were measured by quantitative RT-PCR in neurons derived from three pairs of schizophrenia patients’ *NRXN1^del^* and control iPS cells. Average RQ values (normalized to MAP2 mRNA) were converted to a ratio of control/mutant and plotted on a logarithmic scale. Note that only a limited number of mRNAs were analyzed for pair #2 and only *NNAT* for pair #3 because of its enormous increase in expression in the *NRXN1^del^* neurons. **F**. *NNAT* expression is also increased several-fold in the iPS cells derived from the three *NRXN1^del^* schizophrenia patients analyzed in this study (see B-D for measurement methods). **G**& **H**. *NNAT* expression is not increased in neurons trans-differentiated from genetically engineered stem cells with conditional *NRXN1^del^* mutations. Both neurons derived from previously described ES cells with a conditional *NRXN1* truncation (cTr) that mimics a schizophrenia-associated *NRXN1^del^* mutation (**G**; Pak et al., 2015) and neurons derived from a newly engineered iPS cell line (1215) that causes a conditional deletion of *NRXN1* (**H**; Fig. S4) were analyzed (see B-D for measurement methods). Note that the modest but highly significant increase in *RIMS1* (an active zone protein) expression observed in the patient-derived *NRXN1^del^* neurons is fully reproduced in the engineered lines. Data in C-H are means ± SEM (n ≥ 3 independent cultures for all experiments). Statistical significance was evaluated by Student’s t-tests (* = p<0.05; ** = p<0.01; *** = p<0.001).

Second, we selected test sets of 296 down-regulated and 403 up-regulated DEGs for *NRXN1^del^* vs. control neurons, using a broader set of criteria (uncorrected p<0.05 and |log2(fold-change)|>2). We annotated these DEGs in a Gene Ontology analysis with the ToppFun function of ToppGene (https://toppgene.cchmc.org) (see Methods and Supplementary Data for details). We used hypergeometric tests to compare the fraction of the Test Set (11,880 post-QC genes for this enrichment analysis) that was associated with a given GO term to the fraction of the up- and down-regulated DEGs associated with the same term. There were no significant GO terms for up-regulated DEGs after Bonferroni or FDR correction. For down-regulated DEGs, there were multiple terms in all three GO categories with Bonferroni-corrected significant enrichment of DEGs compared with the Test Set. Characteristic top terms are listed in Fig. 10B (see Supplementary Data for details of significant results and a table showing that the top 9 enrichments are driven by an overlapping set 37 down-regulated DEGs). From a mechanistic standpoint, the upregulated genes notably included *KYAT3*, which encodes kynurenine aminotransferase 3 that is involved in kynurenine metabolism, while the downregulated genes included RELN (involved in neuronal development and signaling) and several transcription factors such as TBX1 (Fig. 10A, S8). The increase in KYAT3 expression was highly reproducible in various experiments, suggesting a true expression change (Fig. S9).

### Quantitative RT-PCR suggests that increased *NNAT* expression is an intrinsic property of patient-derived *NRXN1^del^* iPS cells

In the RNAseq experiments, genes were excluded from patient-control DEG analyses if one or more of the 18 samples had an expression value of ≤ 0.1 in the RNAseq experiments. One of these genes, *NNAT* (encoding neuronatin), demonstrated a striking upregulation in neurons derived from *NRXN1^del^* patients even if the pair of samples with a very low expression value in the control was ignored. Puzzlingly, no upregulation of *NNAT* was observed by RNAseq in engineered *NRXN1^del^* neurons. To explore the possibility that *NRXN1^del^* neurons generated from schizophrenia patients exhibit a specific upregulation of NNAT expression, we performed quantitative RT-PCR experiments on patient-derived *NRXN1^del^* vs. control neurons (Fig. 10C-10E). These measurements confirmed that *NNAT* expression was massively enhanced in patient-derived *NRXN1^del^* neurons. Other neuronal control genes that we tested exhibited no increase except for *RIMS1*, an active zone protein, which exhibited a small but significant increase in expression (Fig. 10C-10E, see Supplementary Data). Although *NNAT* is preferentially produced in neurons, it is also expressed in other cell types. When we measured *NNAT* mRNA levels in patient-derived *NRXN1^del^* vs. control iPS cells, we found also a consistent increase in *NNAT* expression in all patient-derived iPS cells (Fig. 10F). Thus, the increased *NNAT* expression in patient-derived *NRXN1^del^* cells was not specific to neurons.

We next asked whether the increased *NNAT* expression could be a direct consequence of the heterozygous *NRXN1* deletion. We analyzed *NRXN1^del^* neurons obtained from two different engineered, conditionally mutant stem cell lines (see Table 2), our originally described ES cell line (Pak et al., 2015; Fig. 10G) and the newly engineered iPS cell line (Fig. 10H). Both populations of *NRXN1^del^* neurons exhibited no increase in *NNAT* expression, even though they displayed the modest increase in *RIMS1* expression observed in patient-derived *NRXN1^del^* neurons (Fig. 10C, 10D). Thus, *NRXN1^del^* neurons derived from schizophrenia patients exhibited a consistent *NNAT* expression change that was absent from controls and from ‘normal’ neurons in which the same *NRXN1* deletion was engineered, despite the fact that these neurons display the same functional phenotype. A possible reason for the difference in *NNAT* expression between the engineered *NRXN1^del^* and the patient-associated *NRXN1^del^* mutation could be that *NNAT* is an imprinted gene (Kagitani et al., 1997; Evans et al., 2005). Silencing of *NNAT* by imprinting may be maintained in the engineered iPS cell line but not in the patient-derived iPS cell line. An alternative explanation could be that the relative immaturity of the engineered neurons is responsible, but this seems unlikely given that increased *NNAT* expression is already observed in patient-derived *NRXN1^del^* iPS cells, and thus not dependent on neuronal trans-differentiation and maturation.

## DISCUSSION

Schizophrenia is a devastating and prevalent mental disorder with a huge impact on millions of patients and their families. Recent advances in human genetics have identified variations in multiple chromosomal regions and genes that contribute to the genetic risk for schizophrenia and may provide clues to schizophrenia pathogenesis (Coelewij and Curtis, 2018; Kirov, 2015). Enormous progress was made in describing the genetic landscape underlying schizophrenia, resulting in the realization that changes in a large number of genes can contribute to schizophrenia (SCZ working group of PGC, 2014; Pardinas et al., 2018; Marshall et al., 2017; Fromer 2014, Purcell et al., 2014). This genetic heterogeneity supports the notion that schizophrenia is a multifaceted and multidimensional disorder that may not be understandable in terms of a single general pathophysiological mechanism, possibly because no such mechanism exists owing to a great degree of disease heterogeneity, or because many reported findings on mechanisms in schizophrenia were based on underpowered and preliminary experiments.

A large number of 2p16.3 CNVs were described that cause different deletions of chromosomal DNA, but affect expression of only a single gene, *NRXN1* (Marshall et al., 2017, Castronovo et al., 2020). These CNVs strongly predispose to schizophrenia. Although rare, *NRXN1* CNVs affect thousands of patients. As a result, *NRXN1* CNVs are at present the most prevalent known single-gene mutation associated with schizophrenia, although other CNVs involving multiple genes (such as 22q11.2) are more prevalent among schizophrenia patients (Marshall et al., 2017; Hu et al., 2019; Kasem et al., 2018). In addition, *NRXN1^del^* CNVs predispose to other neuropsychiatric and neurodevelopmental disorders (Lowther et al., 2017, Castronovo et al., 2020). Strikingly, our previous study suggested that heterozygous deletions of *NRXN1* in human but not in mouse neurons cause a distinct and robust impairment in neurotransmitter release without changing the development or the morphology of neurons (Pak et al., 2015). This was an exciting finding because it identified a robust phenotype caused by a schizophrenia-associated mutation that could potentially be used to gain further insight into schizophrenia pathophysiology and may even be amenable for drug development. Given the current reproducibility crisis, however, it was necessary to independently assess these results. Thus, the present study aimed for a multidimensional validation of these findings. Specifically, the present study had five objectives: (1) To validate or refute the results obtained with engineered conditionally mutant ES cells, using patient-derived iPS cells with a similar mutation. (2) To confirm that the observed phenotype is independent of a particular laboratory, i.e. that it is replicated at multiple sites. (3) To generate tools and reagents for general use by the scientific community that pursue an understanding of the cellular basis of neuropsychiatric disorders. (4) To determine whether human neurons are uniquely sensitive to a heterozygous loss of *NRXN1* as compared to mouse neurons. (5) To achieve further insights into the mechanisms by which *NRXN1* mutations may predispose to schizophrenia. The present study achieved all of these objectives.

We show that human neurons derived from patients with a heterozygous *NRXN1* deletion (*NRXN1^del^* neurons) exhibit the same phenotype of synaptic impairments as engineered human neurons with a heterozygous *NRXN1* deletion, and that this phenotype is robustly observed with neurons produced and analyzed in different laboratories (Fig. 2–5, 8). This result validates the generality of the synaptic phenotype we described earlier, demonstrating that it is disease-relevant (objectives 1 and 2). Moreover, we engineered a conditional *NRXN1^del^* iPS cell line that is free from commercial restrictions for general use by the scientific community, and that enables production of human neurons with a conditional *NRXN1* deletion and a phenotype identical to what we observed in human patient-derived *NRXN1^del^* neurons and in our original engineered conditional *NRXN1^del^* ES cell line (Fig. 6; objective 3). Furthermore, when we compared human and mouse neurons obtained from stem cells with the same trans-differentiation protocol and carrying essentially the same *NRXN1/Nrxn1* deletion, we observed a robust phenotype only in the human neurons but not the mouse neurons (Fig. 7). Thus a human-specific functional phenotype is produced by the same gene in comparable experimental conditions, suggesting that differences between human and mouse genes may not be solely inherent in the genes themselves, but mediated by the genetic context, which is of potential importance in studying the pathophysiological role of human gene mutations (objective 4). Overall, our results therefore establish that in human neurons, heterozygous *NRXN1* deletions, independent of whether they result from spontaneous unfortunate CNVs or from genetic engineering in stem cells, produce a robust but discrete functional phenotype that consists of a decrease in release probability without a change in synapse numbers of neuronal development. The definition of this synaptic impairment enables a mechanistic approach to a better understanding of schizophrenia pathophysiology.

Besides establishing *NRXN1^del^* neurons as a robust model system and resource for schizophrenia studies, our results provide new information on the pathogenetic mechanisms involved (Fig. 8–10; objective 5). The electrophysiological studies revealed that *NRXN1^del^* neurons exhibit a selective impairment in neurotransmitter release without changes in other neuronal properties, suggesting a circumscribed dysfunction of synaptic transmission that is related to the release machinery. Consistent with this conclusion, we uniformly detected in *NRXN1^del^* neurons an increase in CASK protein levels, in agreement with the notion that a decrease in *NRXN1* signaling via CASK may lead to a compensatory increase in CASK levels (Fig. 8). In the RNAseq studies, few major changes in gene expression were detected, possibly because *NRXN1* is not in itself directly involved in regulating gene expression (Fig. 9, 10). However, several interesting gene expression changes were noted. In all patient-derived *NRXN1^del^* neurons, we detected a significant upregulation in *NNAT* expression. *NNAT’s* exact function is currently unknown, but it appears to perform a regulatory role in secretory tissues, especially the brain (Joseph, 2014; Pitale et al., 2016). We did not detect an upregulation of *NNAT* in *NRXN1^del^* neurons derived from genetically engineered iPS cells, possibly because *NNAT* is an imprinted gene and the imprinting state may differ among iPS cells. The fact that *NNAT* is not increased in the engineered *NRXN1^del^* neurons indicates that the increase in *NNAT* expression is not responsible for the synaptic phenotype that is uniformly observed in the patient-derived and engineered *NRXN1^del^* neurons. Finally, the RNAseq experiments identified a prominent change in the expression of *KYAT3*, a gene linked to schizophrenia pathogenesis that mediates the biosynthesis of kynurenic acid. Kynurenic acid can act as an anticonvulsant, is an antagonist of ionotropic glutamate receptors, and has been proposed to be critically involved in schizophrenia pathogenesis (Erhardt et al., 2007). In particular, kynurenic acid is thought to suppress NMDA-receptor function, which in turn is also regulated by *NRXN1* (Dai et al., 2019) and may be decreased in schizophrenia (Nakazawa and Sapkota, 2020). Thus, an overall picture emerges whereby *NRXN1* dysfunction might operate in the same pathway as kynurenic acid in schizophrenia pathogenesis.

In summary, we here present evidence that the robust synaptic phenotype observed upon *NRXN1* deletions in human neurons is disease-relevant since it is fully reproducible in patient-derived neurons. Moreover, in generating iPS cells with a conditional *NRXN1* deletion we have produced a generally available resource for studying schizophrenia pathophysiology. Finally, we have shown that the heterozygous *NRXN1* deletion phenotype is not observed in mouse neurons under the same experimental conditions, strengthening the rationale for studying disease phenotypes in human neurons. The stage is now set to use these findings and reagents for insight into the molecular mechanisms involved.

## EXPERIMENTAL PROCEDURES

### Patient and control cohort

Case and control donors for this study are described in Table 1. They were drawn from the Molecular Genetics of Schizophrenia (MGS) European-ancestry cohort (Shi et al., 2009; see also information on availability of non-identified clinical information, and biomaterials from the National Institute of Mental Health, nimhgenetics.org, studies 6, 29 and SZ0 for cases, and study 29 for controls; and of genome-wide association study data from dbGAP, accession numbers phs000021.v3.p2 for the “GAIN” and phs000167.v1.p1 for the “NONGAIN” subsets of the genotypes). Cases and controls were age 35-51 at collection. Selection criteria for controls included: polygenic risk score in the bottom 20th percentile based on the Psychiatric Genomics Consortium schizophrenia GWAS data in 2013; employed, but not in a professional/managerial position; married; high school education but without >2 years of college; denied psychiatric or substance dependence disorder based on an online screening online questionnaire (nimhgenetics.org). These individuals are predicted to be at low genetic risk of schizophrenia, and not outliers in cognitive function. For the overall project of which this work was a part, iPS cell lines were derived for 5 schizophrenia cases each with *NRXN1* deletions, 22q11.2 deletions or 16p11.2 duplications, and for 6 controls. Note that aliquots of all subclones of these 21 iPS cell lines are available from NRGR for all 21 individuals, including the six clones used in the present work and the 4 subclones of the engineered conditional *NRXN1* mutation line described below (Table 1).

### GWS and CNV-calling methods

For the six individuals (three *NRXN1* deletion schizophrenia cases and three controls) for whom data are reported in this paper, 33-41x whole-genome DNA sequencing (WGS) was carried out from genomic DNA extracted from whole blood by Macrogen USA (Rockville, MD) on the Illumina X Ten platform, using Trueseq Nano library prep kits for paired-end 150bp reads. Sequencing reads were aligned to the human reference genome (hg19) using BWA (Li et al., 2009). GATK (McKenna et al., 2010) was used to call SNVs and Indels. CNVs were called by MetaSV (Mohiyuddin et al., 2015). Sequencing coverage in bedgraph files was calculated by bedtools genomecov (Quinlan et al., 2010) from bam files. Bedgraph files were then converted to tdf files by igvtools toTDF for visualization in IGV (Robinson et al., 2011).

### iPS cell generation, banking and QC procedures

CD4+ T-cells were obtained from frozen peripheral blood mononuclear cell specimens and reprogrammed using sendai virus method into iPS cells at RUCDR Infinite Biologics. Multiple subclones of iPS cell lines were expanded on mTeSR™1 (Stem Cell Technologies) and Matrigel (BD Biosciences) and passaged using Dispase (Stem Cell Technologies) at RUCDR Infinite Biologics. Upon freezing and banking (~passage 12), pre-freeze mycoplasma testing and morphology testing were performed as well as post-thaw morphology and mycoplasma testing (at RUCDR Infinite Biologics). Other optional tests included karyotype, array CGH, alkaline phosphatase staining, pluritest, and sendai persistence assay. RUCDR frozen cell line ampoules were shipped on liquid nitrogen. At Stanford, iPS cell clones were expanded on mouse embryonic fibroblast feeders in human ES media (DMEM/F12 medium supplemented with 20% KnockOut Serum Replacement, 1x Glutamax, 1x MEM NEAA, 1x sodium pyruvate, 1x Pen/Strep, and 2-Mercaptoethanol (0.1 mM), bFGF (10 ng/mL)) and fed every 2-3 days. Master stock was produced (~10 vials at passage 14-17). Subsequently, working stock was produced (~25 vials at passage 19-21). Clones were checked for karyotype and myclopasma regularly. neurons were differentiated from working stocks and carried on for no more than ~10 passages. NRXN1 cKO iPS cells were maintained as feeder-free cells in mTeSR™1 medium on Matrigel. Cell lines were expanded and banked to create master stock and working stock as mentioned above. All experiments were limited to ~10 passages in culture from working stocks.

### Gene targeting

Previously reported NRXN1 cKO strategy for homologous recombination (Pak et al., 2015) was used to engineer the control C3141a iPS cell line using TALENs (FUJIFILM Cellular Dynamics Inc. engineering team). Multiple subclones were generated and tested for correct genomic integration by PCR and Sanger sequencing. Karyotyping, pluripotency test, mycloplasma test, and post-thaw cell test were performed on the engineered lines. Upon successful engineering, cell lines were shipped on dry ice to multiple sites and viability, genotype, and differentiation were independently tested by individual labs (Stanford and UMass).

### Human *NRXN1* exon 19 PCR genotyping

Genotyping on engineered iPSC line was per-formed using the following primers described previously (Pak et al., 2015): (PCR #1) F: 5’-GATGTGTGCTGCTGTTGCTTTTTGG-3’; R: 5’-AAATGTGCTCAAGAAAGTGGACCAAG-3’; (PCR #2) F: 5’-GGCCTGTGGAAATGTCTGTAACA-3’; R: 5’-TATCAAGGTAC CAGTTGTTTTAAAAGA-3’; (PCR #3) F: 5’-TATTCTGGATCCTTTACTGTTAACTTTT-3’; R: 5’-CTTATCAATGAATTCAGCCCTTGTGGT-3’

### FCDI iN cell engineering

The engineered iPSC lines were generated using piggybac-mediated insertion of the inducible human *Ngn2* expression cassette alongside constitutive rtTet expression. The RUCDR iPSC lines were expanded on Matrigel (Corning) in E8 (Thermo Fisher) with EDTA splitting, and a normal karyotype was confirmed before proceeding. When the iPSCs reached 80% confluence they were dissociated with Accutase (Invitrogen) and resuspended in Ingenio Electroporation Solution (Mirus) with 4 μg of plasmid encoding piggyBac transposase and 4 μg of *Ngn2* plasmid. The *Ngn2* plasmid encoded an rtTET-2A-NeoR cassette expressed by the constitutive EEF1A1 promoter, and an Ngn2-2A-PuroR cassette induced by a Tet responsive promoter, with both elements flanked by piggyBac inverted terminal repeats. Electroporation was performed using the Gene Pulser Xcell (BioRad) with exponential decay at 100 V and 950 μF. The electroporated iPSCs were plated in E8 plus 10 μM Blebbistatin in a Matrigel-coated 6 well plate. The cells were fed with E8 for three or more days, then selected with E8 + 200 μg/ml Geneticin (Gibco) during expansion of an engineered pool up to T-150 scale, followed by cryopreservation. The engineered lines were confirmed to have a normal karyotype before induced neuron generation.

### FCDI iN cell differentiation

Induced neurons were derived from the engineered hiPSC lines according to previous protocol (Zhang et al., 2013) with modifications. In brief, engineered hiPSCs were plated as single cells and cultured in E8 medium (Thermo fisher) for one day. The induction started with medium containing DMEM/F12 (Thermo fisher), N2 (Thermo fisher), MEM non-essential Amino Acids (Thermo fisher), Recombinant Human BDNF (10 ng/ml, R&D Systems), Recombinant Human NT3 (10 ng/ml, R&D Systems) and Doxycycline (5 μM, Clontech) on day 2. Puromycin (1 μg/ml, Thermo fisher) selection was started one day after the induction (day 3). The medium was switched to Neurobasal (Thermo fisher), B27 (Thermo fisher), GlutaMax (Thermo fisher), Recombinant Human BDNF (10 ng/ml, R&D Systems), Recombinant Human NT3 (10 ng/ml, R&D Systems) and Doxycycline (5 μM, Clontech) on day 4. The cells were replated on day 6 with medium containing Cytosine β-D-arabinofuranoside hydrochloride (2.5 μM, Sigma). The cells were harvested and frozen on day 10.

### FCDI iN cell post-thaw culturing

Thawing protocol of frozen CDI neurons is as follows: cryopreserved iN cell vials were transferred from liquid nitrogen thawed in a 37C degrees water bath until a small ice crystal was present. Cells were then added dropwise to 5 ml of room temperature MEM in a 15 ml conical tube, with gentle swirling, and pelleted in a table-top centrifuge preset to 4 degrees at 1000 RPM for 5 minutes. MEM was removed and the cell pellet was resuspended in 1 ml of neuronal culture media (Neurobasal-A medium supplemented with 5% FBS, 2 mM L-Glutamine, 2% B27, 10 μg/l human BDNF, and 10 μg/l human NT-3) and counted using a hemocytometer. Approximately 1.8 million viable cells were recovered from each 1X cryovial. To increase neuronal density for electrophysiological recordings, 200,000 neurons/well were allowed to adhere, in the presence of Y-27632 (10 μM), as a 100 μl bubble placed onto Matrigel-coated glass coverslips harboring a glial cell monolayer. Wells were flooded 1 hour later with neuronal culture medium and media was refreshed every 4-5 days.

### Lentivirus

Lentiviruses were produced as described (Pang et al., 2010, Pak et al., 2015). For all lentiviral vectors, viruses were produced in HEK293T cells (ATCC, VA) by co-transfection with three helper plasmids (pRSV-REV, pMDLg/pRRE and vesicular stomatitis virus G protein expression vector) (10 μg of lentiviral vector DNA and 6 μg of each of the helper plasmid DNA per 75 cm^2^ culture area) using calcium phosphate method (Chen and Okayama, 1987). Lentiviruses were harvested from the medium 48 hr after transfection and either concentrated by high centrifugation for infection at day −1 iN induction or directed harvested for superinfection at day 4 of iN protocol. For concentration of lentiviral particles, viral supernatant were pelleted by high-speed centrifugation (49,000 x g for 90 min), resuspended in MEM, and aliquoted and snap-frozen in liquid N2. Only virus preparations with over 90% infection efficiency as assessed by EGFP expression or puromycin resistance were used for experiments.

### Generation of neurons (human neurons) from iPS cells

Neurons were generated from feeder-dependent iPS cells as described previously (Pak et al., 2018): (Day −1) iPS cells were plated and infected on Matrigel coated 24-well plate. Per 6-well of iPS cells grown on feeders, cells were washed 1x with PBS. 0.5 ml of CTK (0.25% Trypsin no phenol red, 0.1 mg/ml collagenase IV, 0.5 ml 1 mM CaCl2, 20% KnockOut Serum Replacement in sterile water) was added to remove the feeders at 37°C until feeder cells lift off. CTK was removed and washed 2x with PBS. iPS cells were treated with 0.5 ml Accutase briefly, resuspended in 1 ml hES media and spun down at 150 g x 5 min. Cell pellets were resuspended in 50% hES media/50% mTeSR plus Y-27632, and lentiviruses. Typically 200,000 - 300,000 cells were plated per well of a 24-well plate with 1 uL of viruses (rtTA and Ngn2-puro) in 1 ml of mixed media. (Day 0) Half of the virus-containing media was removed and fresh Neural induction medium (NIM; DMEM F12 medium supplemented with 1% N2, 1% non-essential amino acids, 10 μg/l human BDNF, 10 μg/l human NT-3, and 0.2 mg/l mouse laminin) was added supplemented with Doxycycline (2 mg/l) and Y-27632 to induce Ngn2-puro expression. (Day 1-2) 24 hours later, Puromycin (1 mg/l) was added to NIM for 24-48 hours and media was changed daily. (Day 3) neurons were washed 1-2x with PBS. Freshly isolated or previously cryopreserved mouse glial cells (~75,000 cells per well of a 24-well plate) were added in Neural differentiation medium plus Dox (NDM; Neurobasal A medium supplemented with 2% B27, 1% glutamax, 2 μM AraC, 10 μg/l human BDNF, 10 μg/l human NT-3, and 0.2 mg/l mouse laminin). (Day 5) Half of the media was changed using NDM plus Dox. Media change occurred every 2-3 days until Day 10. (Day 10) Half of media was replaced with Maturation medium (MM; MEM supplemented with 0.5% glucose, 0.02% NaHCO3, 0.1 mg/ml transferrin, 5% FBS, 0.5 mM L-glutamine, 2% B27, and 2 μM AraC). Dox was withdrawn from this time point forward. (Day 13) Additional MM (0.5 mL per well of a 24-well plate) was added and neurons were incubated for 7 days without media change. (Day 20) Half of media was replaced with fresh MM. Media change occurred weekly until maturity.

Feeder-independent iPS cells were differentiated as described previously (Pak et al., 2015; Pak et al., 2018): (Day −2) iPS cells were plated on Matrigel coated 6-well plate one day before infection. iPS cells were dissociated with Accutase and plated at cell density of ~5 × 10^4 - 1 × 10^5 cells/well using mTeSR medium with Y-27632. (Day −1) On the day of transduction, media was replaced with 2 mL of mTeSR containing lentiviruses (rtTA and Ngn2-puro). Titer the lentiviruses by adding various amounts of lentiviruses to the cells to achieve complete infectivity and clean selection. (Day 0) After 16-18 hours, virus-containing media was removed and fresh NIM plus Dox was added. (Day 1-2) 24 hours later, Puromycin was added to NIM for 24-48 hours and media was changed daily. (Day 3) neurons (150,000 cells per well of a 24-well plate) together with mouse glial cells (~75,000 cells per well of a 24-well plate) were plated on matrigel coated coverslips in NMD plus Dox. (Day 6) Half of the media was changed using NDM plus Dox. Media change occurred every 2-3 days until Day 10. (Day 10) Half of media was replaced with Maturation medium and Dox was withdrawn from this time point forward. (Day 13) Additional MM (0.5 mL per well of a 24-well plate) was added and neurons were incubated for 7 days without media change. (Day 20) Half of media was replaced with fresh MM. Media change occurred weekly until maturity. Details on lentivirus generation, primary mouse glia isolation, and reagent preparations are published online (Pak et al., 2018).

### RNA isolation and preparation

For bulk RNA-sequencing and qRT-PCR validations, 4 – 4.5 week-old neurons co-cultured with mouse astrocytes were lysed and total RNA was extracted using RNeasy mini kit (Qiagen) according to the manufacturer’s protocol. Purified RNA was further treated with RNase free-DNase (Qiagen) to remove genomic DNA contamination and eluted in 40-60 μL per sample. Concentrations, 260/280, and 260/230 ratios were measured using ND-1000 (Invitrogen).

Bulk RNA-sequencing. RNA-seq libraries were prepared from total RNA from ~4-weeks-old induced human neurons co-cultured with mouse glia (triplicate cultures for all iN cell cultures; duplicate for iPS cell lines), using NEBNext Ultra Directional RNA Library Prep Kit for Illumina (NEB). Paired-end sequencing (2 × 150 base-pair reads) was performed using the Illumina Next-seq 500 platform (Novogene). RNA-seq reads were aligned to human + mouse reference (Ensembl Mus_musculus. GRCm38 and Homo_sapiens. GRCh38 transcriptomes v94) using the stranded library option of Kallisto v0.46.0 (Bray et al., 2016) using a concatenated mouse-human index. Based on comparisons with pure mouse and human RNA samples, the Kallisto pseudo-alignment algorithm was able to accurately distinguish and assign transcripts to the proper species reference. From each Kallisto-processed mouse-human RNAseq sample, an expression matrix was constructed to generate human-specific TPMs (transcripts per million) as follows. Human transcript counts and TPM values were extracted and adjusted based on (1) per million human reads per mouse-human sample; (2) transcript-level TPM values were summed per gene; (3) gene TPM values were converted to log2(TPM+1) values, (4) 19,701 genes were selected with log2(TPM+1) ≥ 1 in at least one of the thirty samples, and (5) the resulting gene-level values were per-sample quantile-polished to reduce sample-sample variability (Zyprych-Walczak et al., 2015). Genes for subsequent analyses were selected from 17,004 gene IDs with log2(TPM+1) ≥ 2 in at least one of the thirty samples. Below we refer to log2(TPM+1) values as “logTPM” values.

Differential Gene Expression Analysis. Prior to DEG analyses, we removed gene types that are prone to mapping errors (fusion genes, antisense RNA genes, overlapping and intronic transcript genes) or incomplete capture by standard library prep (non poly-adenylated short RNA genes, e.g., micro-RNA and small nucleolar genes). For each cell type separately, we determined two values for each gene: max (the highest logTPM value for any culture in the set); and min (the minimum value in the set). Each DEG analysis was restricted to genes with max ≥ 2 and min > 0.1 in the relevant set as described below. The rational for the min criterion was that genes with “near-zero” values ≤ yielded inflated DEG statistics, e.g., in genes with no near-zero values, for 9 patients vs. 9 controls, |log2(fold-change)| > 0.2 or > 0.8 was observed in 15.20% and 0.4%, 60.1% and 6.1% of genes with one near-zero value, and in most cases, that value was inconsistent with the other replicates for the same line, suggesting technical artifacts.

DEG analyses used “moderated” t-tests as implemented in limma v3.42.2 (Law et al., 2014) with the limma-trend procedure (fit = eBayes(fit, trend = TRUE)) that uses an Empirical Bayes Model to effect “shrinkage of the estimated sample variances towards a pooled estimate” (Smyth 2004) which can increase power in the face of culture-culture variability. The use of this approach on logTPM data is conservative, which we consider appropriate with a small cohort. R scripts for DEG analyses are available in Supplementary Material.

DEG and cluster analysis of cell types. To assess the transcriptomic state of the 30 cultures, we carried out a cluster analysis of DEGs contrasting the three cell types and a functional enrichment analysis of these clusters. For 15,000 genes meeting the min/max criteria within one or more culture cell-type subsets (non-engineered neurons, engineered neurons, iPS cells), four limma-trend DEG analyses were carried out (non-engineered, engineered or all neuronal vs. 6 iPS cell cultures; engineered vs. non-engineered neurons), and 7763 DEGs were selected (|t| ≥ 15 in any of the first three analyses and/or |t| ≥ 4 in the fourth). K-means clustering (Pearson correlation distance metric) assigned these DEGs to 11 clusters, which were collapsed to 7 based on cluster similarity (C1-C7, containing 1964, 1011, 525, 394, 261, 279 and 3329 genes respectively). We used ToppFun (ToppGene.cchmc.org, October 5, 2020) (Chen et al., 2009) to determine how many of all 15,000 genes and of the DEGs in each cluster were annotated to each Gene Ontology term. Terms containing at least 10 of the 15,000 genes were retained, and the 13,392 genes mapped to at least one of those terms were considered the Test Set.

For each term, a hypergeometric p-value was computed to determine the probability of the observed or a large number of DEGs being annotated to the term, given the proportion of all 13,392 Test Set genes assigned to that term (see Supplementary Data for full results). The p-values were computed in Excel as (1-hypgeom.dist(DEGhits-1, nDEG,TestSetHits, NTestSet,TRUE), where DEGhits = the number of DEGs assigned to the term, nDEG = the number of DEGs in the Test Set, TestSetHits = the number of Test Set genes assigned to the term, nTestSet = number of genes in the test set, and TRUE computes the cumulative distribution of at most DEGhits-1). This yields the probability of observing N or greater DEGhits in the term, given the expected proportion TestSetHits/nTestSet. This avoids spurious results by using the actual Test Set genes, rather than the entire genome, to determine the expected proportion, because the QC procedure can produce a very different number of genes in a given term compared with the genome. Because the three GO categories (Molecular Function, Biological Process, Cellular Component) are non-independent (they annotate the same genes utilizing available biological information), we adopted a separate 5% alpha level for Bonferroni correction of terms within each category, based on the number of tested terms (MF: 1209, p=4.14E-05; BP: 6405, p=7.81E-06; CC: 846; 5.91E-05).

### Patient vs. control DEGs

The primary DEG analysis contrasted the 9 neuronal cultures from 3 NRXN1^del^/schizophrenia patients vs. 9 cultures from 3 control individuals, restricted to 12,910 post-QC genes that met the min/max criteria in these 18 cultures (the lower N is due to the exclusion of genes that only met min/max criteria in iPS or engineered cells). This included 38 genes that were not in the 15,000 cluster test set because they were not annotated for functional enrichment analyses, but they met QC for the patient-control analyses. Secondary analyses contrasted the 3 replicates for each patient-control pair separately using the the 9×9 Test Set. The 3 paired *NRXN1^del^*/wild-type cultures from the conditional knockout engineered line were analyzed in 12,356 genes (Engineered Test Set) that met min/max criteria in those 6 cultures. These analyses used a limma-trend option to carry out a moderated paired t-test by entering a covariate that coded the 9 deletion-control replicate “pair-groups” (*limma* userGuide, section 9.4.1, https://bioconductor.org/packages/devel/bioc/vignettes/limma/inst/doc/). Analysis commands were: design = model.matrix (~ pair-group + NRXN1-geno); fit = lmFit (data_matrix, design); fit = eBayes (fit, trend = TRUE).

We carried out functional enrichment analysis of 9×9 DEGs (separately for up- and down-regulated DEGs), using the same procedure as described above for cell-type DEGs. The enrichment 9×9 Test Set included 11,781 genes that were annotated to at least one GO term with >10 tested genes. The number of tested terms and Bonferroni-corrected significance thresholds were: MF: 1105 terms, p=5.78E-07; BP: 5962, p=2.33E-09; CC 807, p=3.95E-07.

### qRT-PCR validation

Total RNA (~40 μg per reaction) was reverse-transcribed and PCR amplified using Taqman Fast Virus 1-Step Master Mix (Applied Biosystems) and primetime assays for specific genes (IDT). mRNA levels were quantified by real-time PCR assay using the Quantstudio 3 System and RQ analysis software (Thermo Fisher Scientific). To ensure the specificity of the amplification, titrations of total human brain RNA were used to test primer pairs and only primers that demonstrated a linear amplification were analyzed.

### Primer sequences for qRT-PCR assays

Primetime assay (IDT) sequences are as follows (probe/forward primer/reverse primer): NRXN1: CACCCAAGCCTGAAGAACTGTCCA; GCCATAGGTTTTAGCACTGTTC; CTGTCCCAACATTAAACTTAACTCC GAPDH: CAGCAAGAGCACAAGAGGAAGAGAGA; AGGGTGGTGGACCTCAT; TGAGTGTGGCAGGGACT NNAT: ACATTCTGACATCGCCAGCCGAC; TTTCTTAACCACCCTCCTTCC; AAATCAAAACACCGCACCAG RIMS1: CTCCCAGTCGCACTCCAGGAAAT; CCCAATCCATCAAGAAAATCACT; CGAGCCGAGAGTCTACTGAT MAP2: CACCTGCTGCTTCCTCCACTGT; AGACTTTGTCCTTTGCCTGTT; CCATCACCTGCCTCAGAAC UBE3A: TGACGGTGGCTATACCAGGGACT; CGTTGTAAACTGCAAGAAGAGTC; CCAAGCACTAGAAGAAACTA AGA ATP2A2: CCATCGCCAGTCATAGCTGTAATCTCA; TTTCTTCAGAGCAGGAGCATC; TGAACCCTCCCACAAGTCTA ATP2C1: ACGCTTGTTTGAGGCTGTGGATCT; TTGTCTCACCTGTCAAGCTG; ATACACTTGCCCGAGACTTG TFCP2: CTCTGGCTGGTGGTTTGGTGAAC; CTGAGGTGTGGTTGTTGGTAA; GCAGTTTTTCTCTTGGGGAAG HMX1: CTACCTGAGCAGCGCCGAGC; GAACCAGATCTTAACCTGCGT; GAATCCACCTTCGACCTGAA CAPNS1: TGGCCGTGATGGATAGCGACAC; CCACCTTTTGATGTTGTTCCA; TGATGGTTTTGGCATTGACAC PTPRD: CCTTACCCAAACCTCCAGGAACTCC; ACGTCAGTGTGATGCTTGTAG; TGAAGCAATAGCACAGATCACT UNC50: CTGCTCTGTCGTTTCACTAAATACTTGTTAGAGATG; CATAGCCCCATTCCACATCA; GGTTGTACTCATAGATTGTGTAGGC PARD3: TGCTCGAAGGACTGAAGGTGTGAC; CTACTGCGTCGAACATGAAGA; TTCAGCCTTACCAAGCAACA

### Immunoblotting and protein quantifications

For immunoblotting of proteins from iN cells, neurons were washed 3x with PBS and lysed with RIPA buffer (150 mM NaCl, 5 mM EDTA, 1% Triton X-100, 0.1% SDS, 25 mM Tris-HCl, pH 7.6, and 1x cOmplete protease inhibitor cocktail (Sigma-Aldrich)). Lysates were subjected to SDS-PAGE and immunoblotting using fluorescently labeled secondary antibodies (donkey anti-mouse IRDye 800CW, 1:15,000; donkey anti-rabbit IRDye 800CW, 1:15,000; donkey anti-rabbit IRDye 680, 1:15,000; donkey anti-mouse IRDye 680, 1:15,000; LI-COR Bioscience) and signal detection with an Odyssey Infrared Imager and Odyssey software (LI-COR Biosciences). Signals were normalized for TUJ1 probed on the same blots as loading controls. Antibodies used were as follows: NRXN1 (pan-NRXN-alpha 1:500; G394; TCS), CASK (1:1000; 75-000; Neuromab), MUNC18 (1:500; BD Transduction; 610336), SYT1 (1:1000; 41.1, Synaptic Systems), SYP (1:1000; p580; TCS), SNAP25 (1:1000; p913; TCS), SYB2 (1:1000; CI69-1; TCS), alpha-LIPRIN (1:1000; 4195; TCS), MINT1 (1:500; p730; TCS), VELI (1:500; U049; TCS), PSD95 (1:1000; L667; TCS), GLUA1 (1:500; ab1504; Abcam), NLGN1 (1:1000; 4C12; TCS), CAMKII (1:1000; MAB8699; Milipore), CAMKII-Phospho-286/7 (1:1000; 22B1; Thermo Fisher), TUJ1 (1:1000; 801201; Biolegend).

### Mouse breeding

All mouse work was performed as prescribed by approved protocols at Stanford University. Mice were weaned at 20 d of age and housed in groups of up to five on a 12-h light/dark cycle with food and water ad libitum. Animals were kept in the Stanford Animal Housing Facility with all procedures conforming to the standards set by the National Institutes of Health Guidelines for the Care and Use of Laboratory Mice and approved by the Stanford University Administrative Panel on Laboratory Animal Care.

The following primers were used for genotyping to discriminate between different alleles: Genotyping on Nrxn1 cKO mouse ES cells was performed using following primers: F: 5’-GTAGCCTGTTTACTGCAGTTCATTCC-3’, R:5’-CAAGCACAGGATGTAATGGCCTTTC-3’ (WT: 200bp, floxed alleles: 350bp).

### Mouse ES-iN cells

Nrxn1 conditional KO mouse embryonic stem cells were maintained under the feeder-free conditions in 2i + LIF medium (DMEM/F12: Neurobasal = 1: 1 Medium (Invitrogen), supplemented with 2.5% FBS (Hyclone), N2 (Gibco), B27 (Gibco), Glutamax (Gibco), 1000 U/ml leukemia inhibitory factor (LIF, Millipore), 3mM CHIR99021 (Tocris) and 1 mM PD0325901 (Tocris)). For Neuron Generation, the mES cells were treated with Trypsin-EDTA (0.25%, Gibco); the dissociated cells were infected with lentiviruses containing Ubi-rtTA and TetO-Ngn2-t2a-puromycin, and plated on the Matrigel (BD Bioscience)-coated 6-well plate in 2i + LIF medium on day −4 (1 × 10^5^ cells/well). Medium was changed every day. On day −2, the mES cells were dissociated with Trypsin-EDTA and resuspended in EB medium (DMEM/F12: Neurobasal = 1: 1 Medium (Invitrogen), 10% Knockout Serum Replacement (Gibco), Penicillin/Streptomycin (Gibco), Glutamax (Gibco)) on Petri dish for 2 days. On day 0, the Embryoid Bodies (EBs) were dissociated with Accutase (Innovative Cell Technologies) and infected with Ubi-Cre or Ubi-ΔCre, plated on Matrigel-coated 6-well plate in Neurobasal 1% serum medium (Neurobasal, Glutamax, B27, Pen/Strep, FBS) (2 × 10^5^ cells/well). 2 mg/l Doxycyclin (Sigma) was added to induce the TetO genes expression. On day 1, a 48 hr puromycin selection (2 mg/l, Sigma) period was started. On day 3, the iN cells were dissociated with Accutase and replated with mouse glia cells on Matrigel-coated coverslips in 24-well plate (150k cells/well) in Neurobasal 1% serum medium containing 5 μM 5-fluorodexoyuridine (FUDR, Sigma) and 4 μM Uridine (Sigma) to inhibit glial cell growth. After day 3, 50% of the medium in each well was exchanged every 2 days. FBS (2.5%) was added to the culture medium on day 10 to support glia viability, and iN cells were assayed on day14 in most experiments.

### Immunofluorescence labeling experiments

Cultured iN cells were fixed in 4% paraformaldehyde + 4% sucrose in PBS for 20 min at room temperature, washed three times with 0.2% Triton X-100 in PBS (PBST) for 10 min each wash at room temperature. Cells were incubated with blocking buffer (PBS containing 2% normal goat serum (Sigma-Aldrich) and 0.02% Sodium-azide) for 1 hr at room temperature. Primary antibodies, diluted in blocking buffer, were applied for 1 hour at room temperature then washed 3x with PBST. Secondary antibodies were diluted in blocking buffer and applied for 1 hr at room temperature. Immunolabeled neurons were then mounted on glass slides with Fluoromount-G mounting medium. The following antibodies were used for our analysis: chicken anti-MAP2 (1:1000; ab5392; Abcam) and rabbit anti-synapsin (1:200; E028; TCS). Alexa-546- and Alexa-633-conjugated secondary antibodies were obtained from Invitrogen.

### Image Acquisition and Quantification of Neurite outgrowth and Excitatory Synapses

Image acquisition and quantification were performed as described (Pak et al., 2015). Cells were chosen at random from three or more independent cultures. Images were taken from at least three coverslips per experiment. Fluorescent images were acquired at room temperature with an inverted Nikon A1 Eclipse Ti confocal microscope (Nikon) equipped with a 60 x objective (Apo, NA 1.4) and operated by NIS-Elements AR acquisition software. Images were taken at 1024×1024 pixel resolution with a z-stack distance of 0.5 μm. Laser power and photomutiplier settings were set so bleed-through was negligible between channels. Within the same experiment, the same settings for laser power, PMT gain, and offset were used. These settings provided images where the brightest pixels were just under saturation. General Analysis was performed with NIS-Elements Advanced Research software (Nikon). For quantifications, images were thresholded by intensity to exclude background signals and puncta were quantified by counting the number of puncta whose areas ranged from 0.1-4.0 mm^2^. For each experiment, at least 15 cells per condition were analyzed and the mean and SEM were calculated. Data shown represent the average of the mean values from at least 3 independent experiments. For analysis of dendritic arborizations, neurons were sparsely transfected with AAV-Syn-EGFP to obtain fluorescent images of individual neurons. Images were acquired using Nikon A1 Eclipse Ti confocal microscope (Nikon) equipped with a 20× objective and operated by NIS-Elements AR acquisition software. Images from 20 to 30 neurons per condition (per n=1) were reconstructed using the MetaMorph neurite application, scoring for total neurite length, neurite branch points, and soma area.

For image acquisition and morphometrics performed at Rutgers site, images were taken on a Zeiss LSM700 confocal microscope. The number of Synapsin positive puncta density corresponding to MAP2 dendrites, soma sizes, puncta sizes and dendritic tree area (MAP2) were identified using *Intellicount* (Fantuzzo et al., 2017), a high-throughput, automated synapse quantification program. The system uses adaptive segmentation for selection of synaptic puncta (Synapsin) as well as dendritic boundaries of MAPs signals. The primary processes of dendrites were counted manually after MAP immunostaining. All image quantifications were conducted blindly.

### Electrophysiological recordings

Electrophysiological recordings in cultured iN cells were performed in the whole cell configuration as described previously (Maximov and Südhof, 2005; Zhang et al., 2013; Pak et al., 2015). Patch pipettes were pulled from borosilicate glass capillary tubes (Warner Instruments) using a PC-10 pipette puller (Narishige). The resistance of pipettes filled with intracellular solution varied between 2-4 MOhm. Series resistances were typically maintained in the range of 6-12 MOhm after break-in, and cells with a series resistance of >15 MΩ, or with an unstable series resistance (>20%/min increase) were excluded from further analysis. The bath solution in all experiments contained (in mM): 145 NaCl, 3 KCl, 2 CaCl2, 2 MgCl2, 10 HEPES-NaOH pH 7.4, and 10 glucose; 310 mosm/l.

Intrinsic and action potential (AP) firing properties of iN cells were recorded in current-clamp mode using standard bath solution supplemented with CNQX (20 μM), AP5 (50 μM) and PTX (50 μM) to block all excitatory (AMPAR- and NMDAR-mediated) and inhibitory (GABAAR-mediated) synaptic inputs, respectively. The pipette solution used for current-clamp recordings contained (in mM): 123 K-gluconate, 10 KCl, 8 NaCl, 1 MgCl2, 10 HEPES-KOH pH 7.2, 1 EGTA, 0.1 CaCl2, 1.5 MgATP, Na4GTP and 4 glucose; 295-300 mosm/l. Pipette and series resistances were compensated up to around 80%. In these experiments, first minimal currents were introduced to hold membrane potentials around −70 mV (typically 1-10pA). To elicit action potentials, increasing amount of currents were injected (typically from −10 pA to +60 pA, with 5 pA increments) for 1s in a stepwise manner. Vrest was determined after break-in but before the injection of any currents, as the mean steady-state voltage between any spontaneous sodium spikes. Action potential parameters were determined by analyzing the first AP in any triggered trains of APs. Input resistance (Rin) was calculated as the slope of linear fits of current-voltage plots generated from a series of small current injection steps. To determine whole-cell membrane capacitance (Cm), square wave voltage stimulation was used to produce a pair of decaying exponential current transients that were each analyzed using a least-squares fit technique (Clampfit 9.02).

Action-potential evoked and spontaneous miniature synaptic responses were recorded in voltage-clamp mode with a pipette solution containing (in mM): 135 CsCl, 10 HEPES-CsOH pH 7.2, 5 EGTA, 4 Mg-ATP, 0.3 Na4GTP, and 5 QX-314; 295-300 mosm/l. Presynaptic APs for evoked synaptic responses were triggered by 0.5-ms current (40-90 μA) injections through a local extracellular electrode (FHC concentric bipolar electrode, Catalogue number CBAEC75) placed 100–150μm from the soma of neurons recorded. The frequency, duration, and magnitude of the extracellular stimulus were controlled with a Model 2100 Isolated Pulse Stimulator (A-M Systems, Inc.) synchronized with the Clampex 9 data acquisition software (Molecular Devices). For single AP-evoked synaptic recordings, typically 10-20 stimulation trials were performed with a stimulation interval of 15s at a fixed stimulation intensity and a fixed approximate distance from cellular somas which remained consistent across each neurons and genotypes. To determine paired-pulse ratios (PPRs) of evoked synaptic responses, two sequential APs were elicited with a given time interval. Each paired stimulation intervals (25-5000ms) was repeated at least 3 times with an intertrial-interval of 30s. Stimulation intensities were variable during these recordings to prevent the induction of excessive synaptic network activity following stimulations, as well as to keep the amplitude of evoked synaptic responses <1 nA preventing the saturation of AMPA-type glutamate receptors. Excitatory postsynaptic currents (EPSCs) were pharmacologically isolated with PTX (50 μM) and recorded at a −70mV holding potential, miniature EPSCs (mEPSCs) were monitored in the presence of tetrodotoxin (TTX, 1 μM); all drugs were applied to the bath solution. Spontaneous mEPSCs were analyzed in Clampfit 9.02 (Molecular Devices) using the template matching search and a minimal threshold of 7pA and each event was visually inspected for inclusion or rejection.

Data were digitized at 10 kHz with a 2 kHz low-pass filter using a Multiclamp 700A amplifier (Molecular Devices). Data were analyzed using Clampfit 9.02 software. For all electrophysiological experiments, the experimenter was blind to the condition/genotype of the cultures analyzed. All experiments were performed at room temperature.

### Data and reagent availability

(Note to reviewers: the following arrangements are either completed, or are in process. If this paper is reviewed and we are invited to submit revisions, final accession details will be included.) All biomaterials and data are available to the scientific community as follows:

i. The 3 schizophrenia/*NRXN1^del^* and 3 control iPS cell lines (Table 1) are banked at the Rutgers University Cell and DNA Repository for the NIMH Repository and Genetics (Study 183), and are available to qualified scientists with no commercial restrictions along with additional iPS cell lines for *NRXN1*, 16p11.2^dup^ and 22q11.2^del^ cases and controls (https://www.nimhgenetics.org). RUCDR will bank and share the 4 engineered conditional knockout subclones (Table 2), when COVID-19-related lab restrictions have been lifted.
ii. The NCRCRG Cell Reprogramming Database (CReD) portal (cred-portal.com) will provide open access to dendritic morphometric and synaptic density measurements; protein quantification and qRT-PCR data; electrophysiological tracings, measurements and analyses; PCR validation of the engineered iPS cell line; electrophysiological and morphometric replication data for pairs 1 and 2 (Pang lab); and the Fujifilm-CDI protocol for industrial-scale production of NGN2-induced neurons.
iii. Submission to dbGAP (https://www.ncbi.nlm.nih.gov/gap/) for controlled-access sharing has been approved (project 39338) for the following data: genomic DNA whole genome sequencing data for the 3 *NRXN1^del^* and 3 control individuals studied here and the additional lines noted above; and bulk RNAseq data for all replicates of the non-engineered and engineered pairs presented here.
iv. We are submitting to the NCBI Gene Expression Omnibus (GEO): transcript-level human and mouse read counts; human counts per gene adjusted for transcripts-per-million human reads; the quantile-polished log2(TPM+1) values used in the present analyses, for each of 15,000 final cluster analysis Test Set genes (which include the 12,910 patient-control DEG Test Set genes) for the 30 cultures studied here.
v. Supplementary data files for the present paper are provided as follows (the README sheet in each file provides more detailed information about its contents):

#### 30×15048 logTPM_final.xlsx

Gene IDs and Gene Symbols and quantile polished log2(TPM+1) values for all 15,048 genes that are in the 15,000-gene cluster Test Set, the 12,190-gene patient-control Test Set and/or the 12,356-gene engineered iN Test Set; and columns showing to which sets each gene belongs.

#### 15000_Cluster_TestSet_Info.xlsx

Gene IDs and Symbols; 4 limma t-test values used to select 7763 DEGs that defined clusters 1-7 (Fig. 9B); a column indicating the genes assigned to each cluster; and columns indicating which 12,910 genes were also in the patient-control Test Set and which of these were broadly-defined up- or down-regulated patient-control DEGs.

#### Patient-Control_DEG_results.xlsx

all limma output for patient-control DEG analysis of 12,910 genes in 9 patient vs. 9 control cultures; for separate analyses of each patient-control pair (3 patient vs. 3 control cultures for each pair); and for 12,356 genes for 3 deleted vs. 3 wild-type iN cultures generated from the conditional heterozygous knockout iPS cell line.

#### Enrichment_results_patient-control_and_cluster_DEGs.xlsx

all significant Bonferroni-corrected and 5% FDR GO term results for enrichments analysis of patient-control down-regulated DEGs (no terms were significant for up-regulated DEGs; a table of the down-regulated genes associated with each of the most significant 9 GO terms; and more detailed results for the most significantly enriched GO terms for each of the 7 clusters (Fig. 9B).

## ACKNOWLEDGEMENTS

This work is a component of the National Cooperative Reprogrammed Cell Research Groups (NCRCRG) to Study Mental Illness and was supported by a National Institutes of Health grant to D.F.L., T.C.S. and M.W. (PIs) (5U19MH104172). C.P. received funding from NICHD F32 NRSA postdoctoral fellowship (F32HD078051), Katharine McCormick Advanced Postdoctoral Fellowship from Stanford University, and UMass Amherst faculty startup fund. NIMH Repository received funding from (2U24MH068457). The Stanford Schizophrenia Genetics Research Fund from an anonymous donor provided additional support. We thank Eugenia Jones Ph.D. (Fujifilm Cellular Dynamics Inc., Madison WI) for input and advice, and Steven E. Hyman, M.D. (Broad Institute), Lorenz Studer, M.D. (Sloan Kettering Institute), and Robert Edwards, M.D. (University of California San Francisco) for their assistance throughout the study as members of the Scientific Advisory Board.

Bio-samples for this publication were obtained from NIMH Repository & Genomics Resource, a centralized national biorepository for genetic studies of psychiatric disorders (nimhgenetics.org). These were peripheral blood mononuclear cell samples from study NRGR Study 29. Data and biomaterials generated in Study 29 were collected by the Molecular Genetics of Schizophrenia consortium, part 2 (MGS2), and funded by collaborative NIMH grants to Evanston Northwestern Healthcare/Northwestern University, Evanston, IL, MH59571, Pablo V. Gejman, M.D. (Collaboration Coordinator; PI), Alan R. Sanders, M.D.; Stanford University, Palo Alto, CA, MH61675, Douglas F. Levinson M.D. (PI); Louisiana State University, New Orleans, LA, MH67257, Nancy G. Buccola APRN, B.C., M.S.N. (PI); University of Queensland, Brisbane, Queensland, Australia, MH59588, Bryan J. Mowry, M.D. (PI); University of Colorado, Denver, CO, MH59565, Robert Freedman, M.D. (PI), Ann Olincy, M.D.; Emory University School of Medicine, Atlanta, GA, MH59587, Farooq Amin, M.D. (PI); University of Iowa, Iowa, IA, MH59566, Donald W. Black, M.D. (PI), Raymond R. Crowe, M.D.; Mount Sinai School of Medicine, New York, NY, MH59586, Jeremy M. Silverman, Ph.D. (PI); University of California, San Francisco, CA, MH60870, William F. Byerley, M.D. (PI); Washington University, St. Louis, MO, MH60879, C. Robert Cloninger, M.D. (PI).

## AUTHOR CONTRIBUTIONS

JCM generated and banked all of the iPS cell lines.

JLD contributed to study design, iN cell culture methods and imaging analysis.

BWS led the development of methods for large-scale production of iN cells for the study (managed QC of iPS cell lines and production and QC of iN cells)

JM managed QC of iPS cell lines, production and QC of iN cells, and development of methods for large-scale iN cell production.

MM led the development and generation of the engineered iPS line.

SG, MV assisted with iPSC expansion and iN cell differentiation.

YL generated mouse neurons from ES cells (ES-iN cells).

AH assisted with immunoblotting analysis.

KJ, PD, EB, AM, XZ, and TW performed RNAseq sample processing and data analysis.

CP performed/optimized differentiation of human neurons (iN cells) from patient-derived and engineered iPS cells and from engineered ES cells; contributed to the generation and characterization of engineered iPS cell lines; performed all morphological, protein and RT-PCR experiments; and assisted with RNAseq data analysis.

TD performed all electrophysiological characterization of human neurons derived from iPS cells at Stanford and assisted with morphological imaging analyses.

VM performed all Rutgers site replication experiments of electrophysiology and morphology studies on human neurons (iN cells) generated at Fujifilm-CDI from patient-derived or control iPS cells.

JW performed electrophysiology recordings of neurons derived from engineered human iPS cells and from mouse ES cells.

BJA led the analysis of RNAseq.

ZP oversaw the replication experiments including electrophysiology and image analysis of FCDI-iN cells.

AEU oversaw RNA-seq processing and data analysis

CP, DFL, MW, and TCS planned the experiments, analyzed the data, and wrote the paper in consultation with all co-authors.

## REFERENCES

Abbott, L.F., and Regehr, W.G. (2004). Synaptic computation. Nature 431, 796–803.

Ahn, K., Gotay, N., Andersen, T.M., Anvari, A.A., Gochman, P., Lee, Y., Sanders, S., Guha, S., Darvasi, A., Glessner, J.T., et al. (2014). High rate of disease-related copy number variations in childhood onset schizophrenia. Mol. Psychiatry 19, 568–572.

Anderson, G.R., Aoto, J., Tabuchi, K., Földy, C., Covy, J., Yee, A.X., Wu, D., Lee, S.-J., Chen, L., Malenka, R.C., et al. (2015). Β-neurexins control neural circuits by regulating synaptic endocannabinoid signaling. Cell 162, 593–606.

Aoto, J., Martinelli, D.C., Malenka, R.C., Tabuchi, K., and Südhof, T.C. (2013). Presynaptic neurexin-3 alternative splicing trans-synaptically controls postsynaptic AMPA receptor trafficking. Cell 154, 75–88.

Aoto, J., Földy, C., Ilcus, S.M.C., Tabuchi, K., and Südhof, T.C. (2015). Distinct circuit-dependent functions of presynaptic neurexin-3 at GABAergic and glutamatergic synapses. Nat. Neurosci. 18, 997–1007.

Bray, N.L., Pimentel, H., Melsted, P., and Pachter, L. (2016). Near-optimal probabilistic RNA-seq quantification. Nat. Biotechnol. 34, 525–527.

Butz, S., Okamoto, M., and Südhof, T.C. (1998) A tripartite protein complex with the potential to couple synaptic vesicle exocytosis to cell adhesion in brain. Cell 94, 773–782.

Castronovo, P., Baccarin, M., Ricciardello, A., Picinelli, C., Tomaiuolo, P., Cucinotta, F., Frittoli, M., Lintas, C., Sacco, R., and Persico, A.M. (2020). Phenotypic spectrum of NRXN1 mono- and bi-allelic deficiency: A systematic review. Clin. Genet. 97, 125–137.

Chen, C., and Okayama, H. (1987). High-efficiency transformation of mammalian cells by plasmid DNA. Mol. Cell. Biol. 7, 2745–2752.

Chen, J., Bardes, E.E., Aronow, B.J., and Jegga, A.G. (2009). ToppGene Suite for gene list enrichment analysis and candidate gene prioritization. Nucleic Acids Res. 37, W305–311.

Chen, L.Y., Jiang, M., Zhang, B., Gokce, O., and Südhof, T.C. (2017). Conditional Deletion of All Neurexins Defines Diversity of Essential Synaptic Organizer Functions for Neurexins. Neuron 94, 611–625.e4.

Ching, M.S.L., Shen, Y., Tan, W.-H., Jeste, S.S., Morrow, E.M., Chen, X., Mukaddes, N.M., Yoo, S.-Y., Hanson, E., Hundley, R., et al. (2010). Deletions of NRXN1 (neurexin-1) predispose to a wide spectrum of developmental disorders. Am. J. Med. Genet. B Neuropsychiatr. Genet. 153B, 937–947.

Coelewij, L., and Curtis, D. (2018). Mini-review: Update on the genetics of schizophrenia. Ann. Hum. Genet. 82, 239–243.

Cohen AR, Woods DF, Marfatia SM, Walther Z, Chishti AH, Anderson JM. Human CASK/LIN-2 binds syndecan-2 and protein 4.1 and localizes to the basolateral membrane of epithelial cells. J Cell Biol. 1998 Jul 13;142(1):129–38.

Dai, J., Aoto, J., and Südhof, T.C. (2019). Alternative splicing of presynaptic neurexins differentially controls postsynaptic nmda and ampa receptor responses. Neuron 102, 993–1008.e5.

Erhardt, S., Schwieler, L., Nilsson, L., Linderholm, K., and Engberg, G. (2007). The kynurenic acid hypothesis of schizophrenia. Physiol. Behav. 92, 203–209.

Evans, H.K., Weidman, J.R., Cowley, D.O., and Jirtle, R.L. (2005). Comparative phylogenetic analysis of blcap/nnat reveals eutherian-specific imprinted gene. Mol. Biol. Evol. 22, 1740–1748.

Fantuzzo, J.A., Mirabella, V.R., Hamod, A.H., Hart, R.P., Zahn, J.D., and Pang, Z.P. (2017). Intellicount: High-Throughput Quantification of Fluorescent Synaptic Protein Puncta by Machine Learning. ENeuro 4.

Flaherty, E., Zhu, S., Barretto, N., Cheng, E., Deans, P.J.M., Fernando, M.B., Schrode, N., Francoeur, N., Antoine, A., Alganem, K., et al. (2019). Neuronal impact of patient-specific aberrant NRXN1α splicing. Nat. Genet. 51, 1679–1690.

Fromer, M., Pocklington, A.J., Kavanagh, D.H., Williams, H.J., Dwyer, S., Gormley, P., Georgieva, L., Rees, E., Palta, P., Ruderfer, D.M., et al. (2014). De novo mutations in schizophrenia implicate synaptic networks. Nature 506, 179–184.

Fromer, M., Roussos, P., Sieberts, S.K., Johnson, J.S., Kavanagh, D.H., Perumal, T.M., Ruderfer, D.M., Oh, E.C., Topol, A., Shah, H.R., et al. (2016). Gene expression elucidates functional impact of polygenic risk for schizophrenia. Nat. Neurosci. 19, 1442–1453.

Guilmatre, A., Dubourg, C., Mosca, A.-L., Legallic, S., Goldenberg, A., Drouin-Garraud, V., Layet, V., Rosier, A., Briault, S., Bonnet-Brilhault, F., et al. (2009). Recurrent rearrangements in synaptic and neurodevelopmental genes and shared biologic pathways in schizophrenia, autism, and mental retardation. Arch. Gen. Psychiatry 66, 947–956.

Hackett, A., Tarpey, P.S., Licata, A., Cox, J., Whibley, A., Boyle, J., Rogers, C., Grigg, J., Partington, M., Stevenson, R.E., et al. (2010). CASK mutations are frequent in males and cause X-linked nystagmus and variable XLMR phenotypes. Eur. J. Hum. Genet. 18, 544–552.

Hall, L.S., Medway, C.W., Pain, O., Pardiñas, A.F., Rees, E.G., Escott-Price, V., Pocklington, A., Bray, N.J., Holmans, P.A., Walters, J.T.R., et al. (2020). A transcriptome-wide association study implicates specific pre- and post-synaptic abnormalities in schizophrenia. Hum. Mol. Genet. 29, 159–167.

Hefft, S., Kraushaar, U., Geiger, J.R.P., and Jonas, P. (2002). Presynaptic short-term depression is maintained during regulation of transmitter release at a GABAergic synapse in rat hippocampus. J. Physiol. (Lond.) 539, 201–208.

Hsueh YP, Yang FC, Kharazia V, Naisbitt S, Cohen AR, Weinberg RJ, Sheng M Direct interaction of CASK/LIN-2 and syndecan heparan sulfate proteoglycan and their overlapping distribution in neuronal synapses. J Cell Biol. 1998 Jul 13;142(1):139–51.

Hu, Z., Xiao, X., Zhang, Z., and Li, M. (2019). Genetic insights and neurobiological implications from NRXN1 in neuropsychiatric disorders. Mol. Psychiatry 24, 1400–1414.

Huang, A.Y., Yu, D., Davis, L.K., Sul, J.H., Tsetsos, F., Ramensky, V., Zelaya, I., Ramos, E.M., Osiecki, L., Chen, J.A., et al. (2017). Rare Copy Number Variants in NRXN1 and CNTN6 Increase Risk for Tourette Syndrome. Neuron 94, 1101–1111.e7.

Hyman, S.E. (2015). Enlisting hESCs to Interrogate Genetic Variants Associated with Neuropsychiatric Disorders. Cell Stem Cell 17, 253–254.

Joseph, R.M. (2014). Neuronatin gene: Imprinted and misfolded: Studies in Lafora disease, diabetes and cancer may implicate NNAT-aggregates as a common downstream participant in neuronal loss. Genomics 103, 183–188.

Kagitani, F., Kuroiwa, Y., Wakana, S., Shiroishi, T., Miyoshi, N., Kobayashi, S., Nishida, M., Kohda, T., Kaneko-Ishino, T., and Ishino, F. (1997). Peg5/Neuronatin is an imprinted gene located on sub-distal chromosome 2 in the mouse. Nucleic Acids Res. 25, 3428–3432.

Kasem, E., Kurihara, T., and Tabuchi, K. (2018). Neurexins and neuropsychiatric disorders. Neurosci. Res. 127, 53–60.

Kirov, G., Rees, E., Walters, J.T.R., Escott-Price, V., Georgieva, L., Richards, A.L., Chambert, K.D., Davies, G., Legge, S.E., Moran, J.L., et al. (2014). The penetrance of copy number variations for schizophrenia and developmental delay. Biol. Psychiatry 75, 378–385.

Kirov, G. (2015). CNVs in neuropsychiatric disorders. Hum. Mol. Genet. 24, R45–49.

Law, C.W., Chen, Y., Shi, W., and Smyth, G.K. (2014). voom: Precision weights unlock linear model analysis tools for RNA-seq read counts. Genome Biology 15, R29.

Li, H., and Durbin, R. (2009). Fast and accurate short read alignment with Burrows-Wheeler transform. Bioinformatics 25, 1754–1760.

Liu, Y., Hu, Z., Xun, G., Peng, Y., Lu, L., Xu, X., Xiong, Z., Xia, L., Liu, D., Li, W., et al. (2012). Mutation analysis of the NRXN1 gene in a Chinese autism cohort. J Psychiatr Res 46, 630–634.

Luo, F., Sclip, A., Jiang, M., and Südhof, T.C. (2020). Neurexins cluster Ca2+ channels within the presynaptic active zone. EMBO J. 39, e103208.

Lowther, C., Speevak, M., Armour, C.M., Goh, E.S., Graham, G.E., Li, C., Zeesman, S., Nowaczyk, M.J.M., Schultz, L.-A., Morra, A., et al. (2017). Molecular characterization of NRXN1 deletions from 19, 263 clinical microarray cases identifies exons important for neurodevelopmental disease expression. Genet Med 19, 53–61.

Marshall, C.R., Howrigan, D.P., Merico, D., Thiruvahindrapuram, B., Wu, W., Greer, D.S., Antaki, D., Shetty, A., Holmans, P.A., Pinto, D., et al. (2017). Contribution of copy number variants to schizophrenia from a genome-wide study of 41,321 subjects. Nat. Genet. 49, 27–35.

Malhotra, D., and Sebat, J. (2012). Cnvs: harbingers of a rare variant revolution in psychiatric genetics. Cell 148, 1223–1241.

Maximov, A., Pang, Z.P., Tervo, D.G.R., and Südhof, T.C. (2007). Monitoring synaptic transmission in primary neuronal cultures using local extracellular stimulation. J. Neurosci. Methods 161, 75–87.

Missler, M., Zhang, W., Rohlmann, A., Kattenstroth, G., Hammer, R.E., Gottmann, K., and Südhof, T.C. (2003). Alpha-neurexins couple Ca2+ channels to synaptic vesicle exocytosis. Nature 423, 939–948

McKenna, A., Hanna, M., Banks, E., Sivachenko, A., Cibulskis, K., Kernytsky, A., Garimella, K., Altshuler, D., Gabriel, S., Daly, M., et al. (2010). The Genome Analysis Toolkit: a MapReduce framework for analyzing next-generation DNA sequencing data. Genome Res. 20, 1297–1303.

Mohiyuddin, M., Mu, J.C., Li, J., Bani Asadi, N., Gerstein, M.B., Abyzov, A., Wong, W.H., and Lam, H.Y.K. (2015). MetaSV: an accurate and integrative structural-variant caller for next generation sequencing. Bioinformatics 31, 2741–2744.

Nag, A., Bochukova, E.G., Kremeyer, B., Campbell, D.D., Muller, H., Valencia-Duarte, A.V., Cardona, J., Rivas, I.C., Mesa, S.C., Cuartas, M., et al. (2013). CNV analysis in Tourette syndrome implicates large genomic rearrangements in COL8A1 and NRXN1. PLoS ONE 8, e59061.

Najm, J., Horn, D., Wimplinger, I., Golden, J.A., Chizhikov, V.V., Sudi, J., Christian, S.L., Ullmann, R., Kuechler, A., Haas, C.A., et al. (2008). Mutations of CASK cause an X-linked brain malformation phenotype with microcephaly and hypoplasia of the brainstem and cerebellum. Nat. Genet. 40, 1065–1067.

Nakazawa, K., and Sapkota, K. (2020). The origin of NMDA receptor hypofunction in schizophrenia. Pharmacol. Ther. 205, 107426.

Neale, B.M., Kou, Y., Liu, L., Ma’ayan, A., Samocha, K.E., Sabo, A., Lin, C.-F., Stevens, C., Wang, L.-S., Makarov, V., et al. (2012). Patterns and rates of exonic de novo mutations in autism spectrum disorders. Nature 485, 242–245.

Neher, E., and Brose, N. (2018). Dynamically Primed Synaptic Vesicle States: Key to Understand Synaptic Short-Term Plasticity. Neuron 100, 1283–1291.

Pardiñas, A.F., Holmans, P., Pocklington, A.J., Escott-Price, V., Ripke, S., Carrera, N., Legge, S.E., Bishop, S., Cameron, D., Hamshere, M.L., et al. (2018). Common schizophrenia alleles are enriched in mutation-intolerant genes and in regions under strong background selection. Nat. Genet. 50, 381–389.

Pak, C., Danko, T., Zhang, Y., Aoto, J., Anderson, G., Maxeiner, S., Yi, F., Wernig, M., and Südhof, T.C. (2015). Human neuropsychiatric disease modeling using conditional deletion reveals synaptic transmission defects caused by heterozygous mutations in nrxn1. Cell Stem Cell 17, 316–328.

Pak, C., Grieder, S., Yang, N., Zhang, Y., Wernig, M., and Sudhof, T. (2018). Rapid generation of functional and homogeneous excitatory human forebrain neurons using Neurogenin-2 (Ngn2). Protocol Exchange. 10.1038/protex.2018.082

Panchision, D.M. (2016). Concise Review: Progress and Challenges in Using Human Stem Cells for Biological and Therapeutics Discovery: Neuropsychiatric Disorders. Stem Cells 34, 523–536.

Pang, Z.P., Yang, N., Vierbuchen, T., Ostermeier, A., Fuentes, D.R., Yang, T.Q., Citri, A., Sebastiano, V., Marro, S., Südhof, T.C., et al. (2011). Induction of human neuronal cells by defined transcription factors. Nature 476, 220–223.

Piluso, G., Carella, M., D’Avanzo, M., Santinelli, R., Carrano, E.M., D’Avanzo, A., D’Adamo, A.P., Gasparini, P., and Nigro, V. (2003). Genetic heterogeneity of FG syndrome: a fourth locus (Fgs4) maps to Xp11.4-p11.3 in an Italian family. Hum. Genet. 112, 124–130.

Pitale, P.M., Howse, W., and Gorbatyuk, M. (2017). Neuronatin Protein in Health and Disease. J. Cell. Physiol. 232, 477–481.

Priyamvada M Pitale 1, Wayne Howse 1, Marina Gorbatyuk 1 Neuronatin Protein in Health and Disease J Cell Physiol 2017 Mar;232(3):477–481.

Purcell, S.M., Moran, J.L., Fromer, M., Ruderfer, D., Solovieff, N., Roussos, P., O’Dushlaine, C., Chambert, K., Bergen, S.E., Kähler, A., et al. (2014). A polygenic burden of rare disruptive mutations in schizophrenia. Nature 506, 185–190.

Quinlan, A.R., and Hall, I.M. (2010). BEDTools: a flexible suite of utilities for comparing genomic features. Bioinformatics 26, 841–842.

Rees, E., Walters, J.T.R., Georgieva, L., Isles, A.R., Chambert, K.D., Richards, A.L., Mahoney-Davies, G., Legge, S.E., Moran, J.L., McCarroll, S.A., et al. (2014). Analysis of copy number variations at 15 schizophrenia-associated loci. Br J Psychiatry 204, 108–114.

Ripke S, Group SW, O’Donovan M. (2017). Current Status of Schizophrenia Gwas. European Neuropsychopharmacology. 27(Supplement 3):S415.

Robinson, J.T., Thorvaldsdóttir, H., Winckler, W., Guttman, M., Lander, E.S., Getz, G., and Mesirov, J.P. (2011). Integrative genomics viewer. Nat. Biotechnol. 29, 24–26.

Saitsu, H., Kato, M., Osaka, H., Moriyama, N., Horita, H., Nishiyama, K., Yoneda, Y., Kondo, Y., Tsurusaki, Y., Doi, H., et al. (2012). CASK aberrations in male patients with Ohtahara syndrome and cerebellar hypoplasia. Epilepsia 53, 1441–1449.

Sanders, S.J., Murtha, M.T., Gupta, A.R., Murdoch, J.D., Raubeson, M.J., Willsey, A.J., Ercan-Sencicek, A.G., DiLullo, N.M., Parikshak, N.N., Stein, J.L., et al. (2012). De novo mutations revealed by whole-exome sequencing are strongly associated with autism. Nature 485, 237–241.

Schizophrenia Working Group of the Psychiatric Genomics Consortium (2014). Biological insights from 108 schizophrenia-associated genetic loci. Nature 511, 421–427.

Sebat, J., Lakshmi, B., Troge, J., Alexander, J., Young, J., Lundin, P., Månér, S., Massa, H., Walker, M., Chi, M., et al. (2004). Large-scale copy number polymorphism in the human genome. Science 305, 525–528.

Sebat, J., Lakshmi, B., Malhotra, D., Troge, J., Lese-Martin, C., Walsh, T., Yamrom, B., Yoon, S., Krasnitz, A., Kendall, J., et al. (2007). Strong association of de novo copy number mutations with autism. Science 316, 445–449.

Shi, J., Levinson, D.F., Duan, J., Sanders, A.R., Zheng, Y., Pe’er, I., Dudbridge, F., Holmans, P.A., Whittemore, A.S., Mowry, B.J., et al. (2009). Common variants on chromosome 6p22.1 are associated with schizophrenia. Nature 460, 753–757.

Smyth, G.K. (2004). Linear models and empirical bayes methods for assessing differential expression in microarray experiments. Statistical Applications in Genetics and Molecular Biology 3, Article3.

Südhof, T.C. (2017). Synaptic neurexin complexes: a molecular code for the logic of neural circuits. Cell 171, 745–769.

Sullivan, P.F., and Geschwind, D.H. (2019). Defining the Genetic, Genomic, Cellular, and Diagnostic Architectures of Psychiatric Disorders. Cell 177, 162–183.

Szatmari, P., Autism Genome Project Consortium, Paterson, A.D., Zwaigenbaum, L., Roberts, W., Brian, J., Liu, X.-Q., Vincent, J.B., Skaug, J.L., Thompson, A.P., et al. (2007). Mapping autism risk loci using genetic linkage and chromosomal rearrangements. Nat. Genet. 39, 319–328.

Tabuchi, K., Biederer, T., and Südhof, T.C. (2002) CASK participates in alternative tripartite complexes in which Mint 1 competes for binding with Caskin 1, a novel CASK binding protein. J. Neurosci. 22, 4264–4273.

Thamban, T., Sowpati, D.T., Pai, V., Nithianandam, V., Abe, T., Shioi, G., Mishra, R.K., and Khosla, S. (2019). The putative Neuronatin imprint control region is an enhancer that also regulates the Blcap gene. Epigenomics 11, 251–266.

Trotter, J.H., Hao, J., Maxeiner, S., Tsetsenis, T., Liu, Z., Zhuang, X., and Südhof, T.C. (2019). Synaptic neurexin-1 assembles into dynamically regulated active zone nanoclusters. J. Cell Biol. 218, 2677–2698.

Vierbuchen, T., Ostermeier, A., Pang, Z.P., Kokubu, Y., Südhof, T.C., and Wernig, M. (2010). Direct conversion of fibroblasts to functional neurons by defined factors. Nature 463, 1035–1041.

Wei Z, Zheng S, Spangler SA, Yu C, Hoogenraad CC, Zhang M. Liprin-mediated large signaling complex organization revealed by the liprin-α/CASK and liprin-α/liprin-β complex structures. Mol Cell. 2011 Aug 19;43(4):586–98.

Wapinski, O.L., Vierbuchen, T., Qu, K., Lee, Q.Y., Chanda, S., Fuentes, D.R., Giresi, P.G., Ng, Y.H., Marro, S., Neff, N.F., et al. (2013). Hierarchical mechanisms for direct reprogramming of fibroblasts to neurons. Cell 155, 621–635.

Yi, F., Danko, T., Botelho, S.C., Patzke, C., Pak, C., Wernig, M., and Südhof, T.C. (2016). Autism-associated SHANK3 haploinsufficiency causes Ih channelopathy in human neurons. Science 352, aaf2669.

Zhang, Y., Pak, C., Han, Y., Ahlenius, H., Zhang, Z., Chanda, S., Marro, S., Patzke, C., Acuna, C., Covy, J., et al. (2013). Rapid single-step induction of functional neurons from human pluripotent stem cells. Neuron 78, 785–798.

Zyprych-Walczak, J., Szabelska, A., Handschuh, L., Górczak, K., Klamecka, K., Figlerowicz, M., and Siatkowski, I. (2015). The Impact of Normalization Methods on RNA-Seq Data Analysis. BioMed Research International 2015, 621690.

