## Supplementary Figures and Legends for "Cross-Platform Validation of Neurotransmitter Release Impairments in Schizophrenia Patient-Derived *NRXN1*-Mutant Neurons"

### SUPPLEMENTARY FIGURES and FIGURE LEGENDS

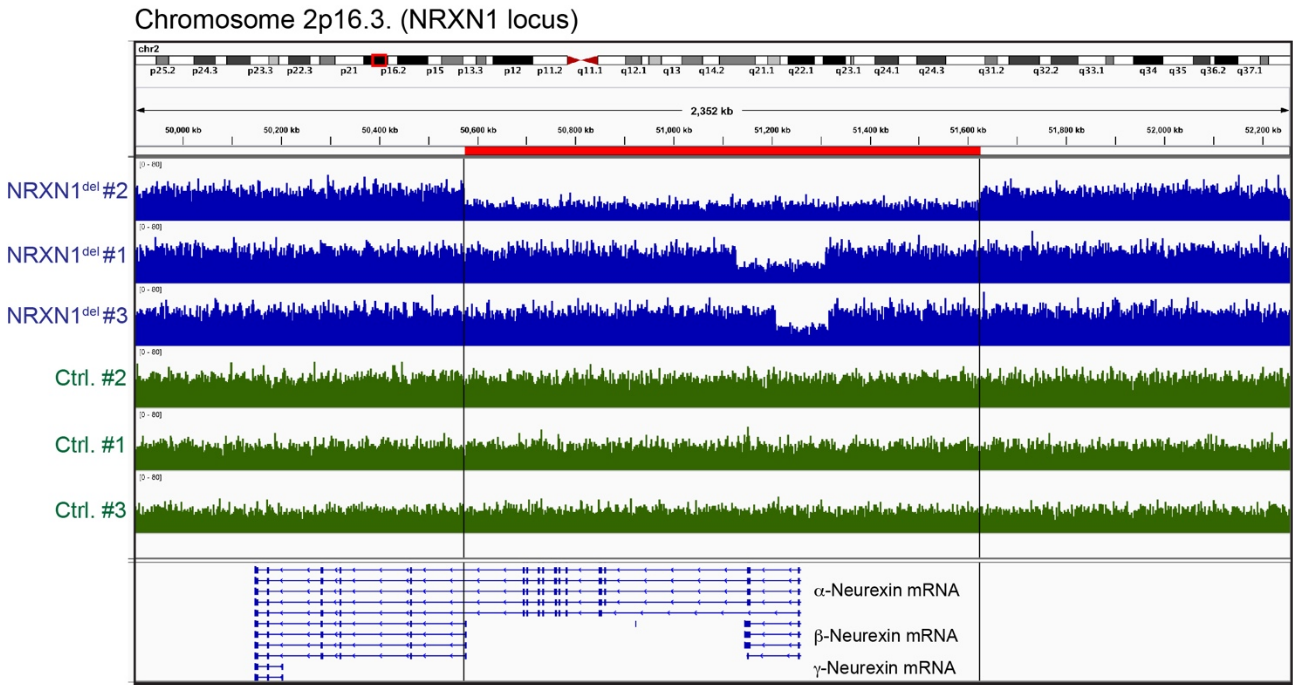


Figure S1: **Copy-number-variation (CNV) plots of the 2p16.3 region from control individuals and patients with *NRXN1* deletions**

Copy-number-variation (CNV) plots showing the *NRXN1* genomic region at 2p16.3 (based on hg19). Blue plots represent schizophrenia $NRXN1^{del}$ patients ($NRXN1^{del}$) and green plots represent control individuals (Ctrl.). Sizes of deletions are described in Table 1 ($NRXN1^{del}$ #1: 0.18 mb; $NRXN1^{del}$ #2: 1.05 mb; $NRXN1^{del}$ #3: 0.10 mb). The locations of *NRXN1* transcript variants are depicted below.

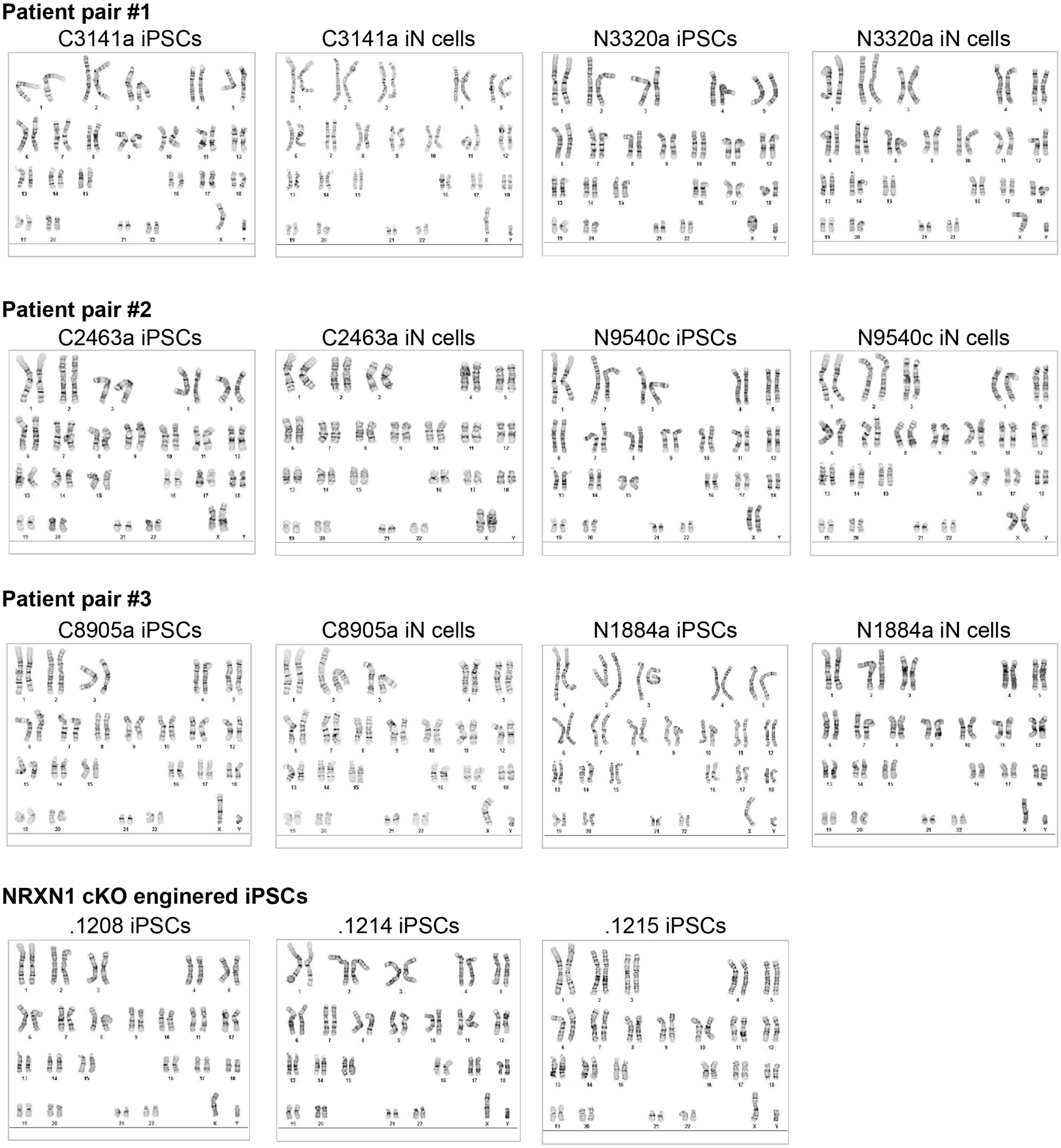


Figure S2: **Karyotyping results for human iPS cell lines and neurons (iN cells)**

Chromosomal analysis (WiCell) validates normal karyotypes of iPS cell lines and neurons generated for each line as indicated. No clonal abnormalities were detected at the stated level of G-band resolution (400-550). Analyses were carried out at FCDI.

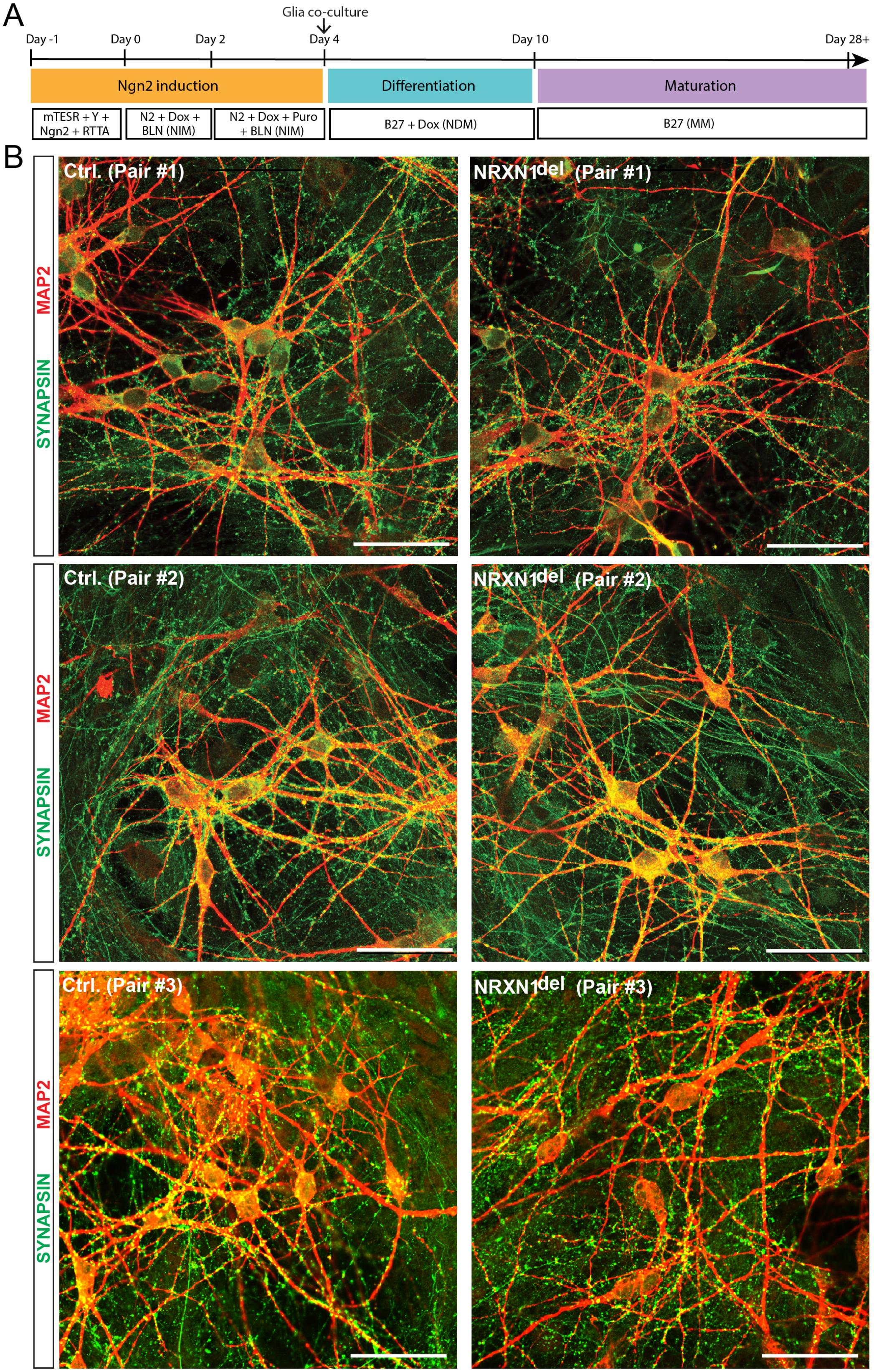


Figure S3: **Generation of neurons from pairs of iPS cells: Protocol (A) and morphology of control and patient-derived *NRXN1*$^{del}$ neurons (B)**

**A**. Schematic of the experimental flow for the Ngn2-induced trans-differentiation of feeder-dependent iPS cells into neurons (abbreviations used: NIM, neural induction medium; NDM, neural differentiation medium; MM, maturation medium; B, BDNF; L, laminin; N, NT3).

**B**. Representative confocal images of Ngn2-induced neurons generated from Ctrl. and $NRXN1^{del}$ pairs. Neurons were immunostained for MAP2 (dendrites; red) and Synapsin-1 (pre-synaptic markers; green) at 4 weeks in culture (scale bar = 50 μm).

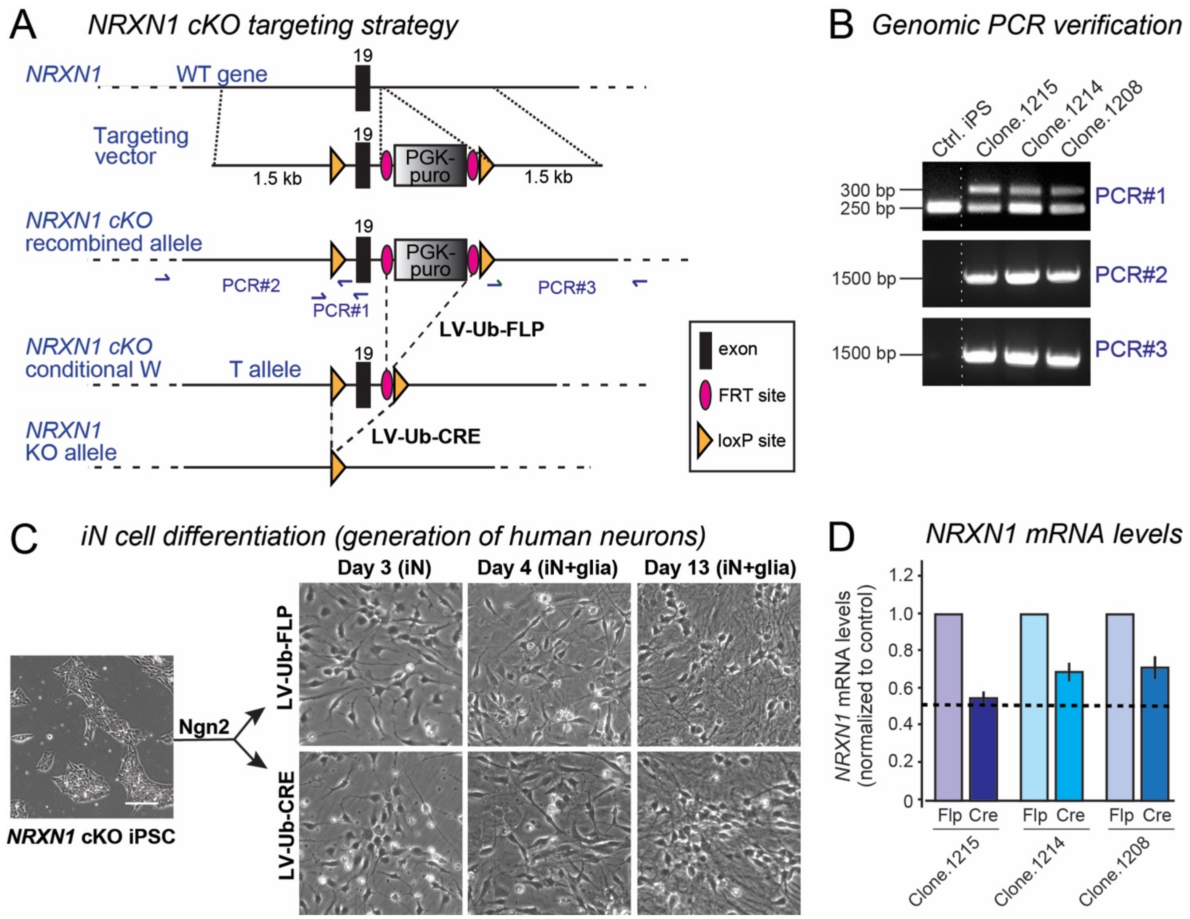


Figure S4: **Generation of a newly engineered line of iPS cells with a conditional heterozygous *NRXN1* deletion**

**A**. *NRXN1* cKO targeting strategy that replicates the approach used previously in H1 ES cells (Pak et al., 2015) and in mice (Chen et al., 2017; Trotter et al., 2019). Using TALEN-assisted homologous recombination, the first exon shared by *NRXN1*$\alpha$ and *NRXN1*$\beta$ isoforms (exon 19; Tabuchi and Südhof, 2001) was flanked with LoxP sites, and an FRT-flanked PGK-puromycin resistance cassette (PGK-puro) was inserted into the intron following exon 19. Flp-recombinase deletes PGK-puro, creating a wild-type allele, whereas Cre-recombinase deletes the exon 19, resulting in a KO allele. The conditional *NRXN1* mutation was introduced into the control iPS cell line C3141a and validated by PCR, sequencing, and mRNA measurements upon FLP and CRE treatments in induced neurons.

**B**. PCR validation of the correct integration of the targeting construct using primers outlined in A. PCR#1 confirms the presence of both wild-type (lower bands) and targeted alleles (upper bands) for the *NRXN1* cKO. PCR#2-3 detect only the targeted cKO allele due to the partially annealing sequences of primers to the loxP sites in combination with primers that are present outside the homology arms.

**C**. Workflow of trans-differentiation of engineered *NRXN1* cKO iPS cells into induced neurons. Upon Ngn2 induction, Flp- or Cre-recombinases are separately introduced via lentiviral infection. Morphologies of induced neurons at developmental ages before and after mouse glia addition are shown (scale bar, 100 $\mu$m).

**D**. Quantitative RT-PCR measurements confirms downregulation of *NRXN1* mRNA in Cre-treated compared to Flp-treated induced neurons. Average ∆Ct values (normalized to MAP2 mRNA) were converted to a ratio of Flp/Cre. Data are means ± SEM (n=4-5- independent cultures). Statistical analyses were performed by Student's *t* test comparing test samples to the control (*$p$<0.05).

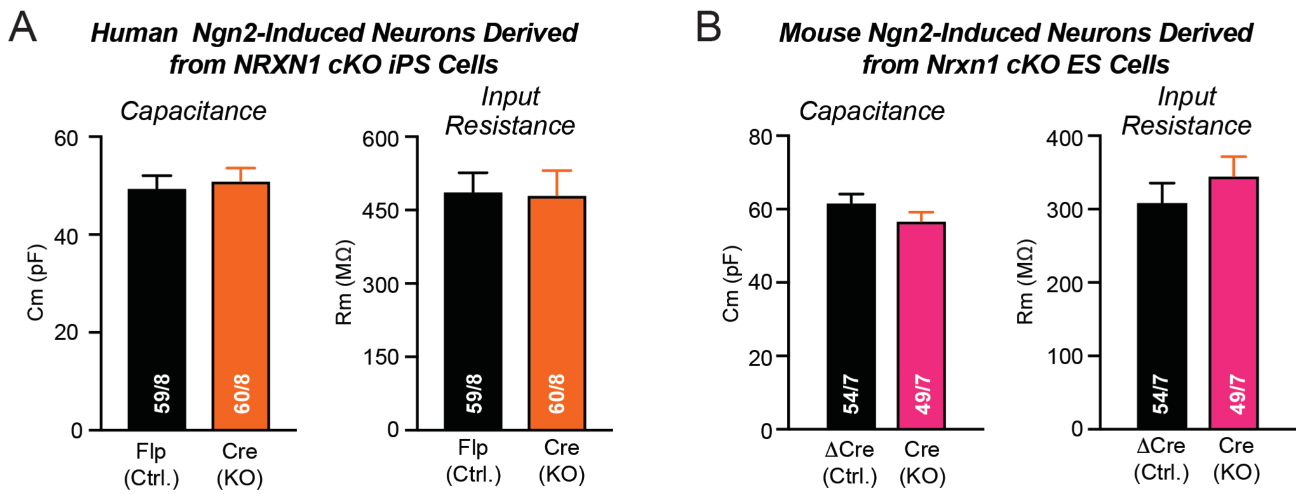


Figure S5: **Normal capacitance and input resistance in human neurons trans-differentiated from the newly engineered *NRXN1* cKO iPS cell line (1215sub) (A) and in mouse iN cells with a heterozygous deletion of *Nrxn1* trans-differentiated from the newly generated mouse *Nrxn1* cKO ES cell line**

**A**. Capacitance and input resistance of human control (Flp) and *NRXN1* heterozygous KO neurons (Cre) trans-differentiated from the newly engineered iPS cell line.

**B**. Capacitance and input resistance of mouse control (ΔCre) and *Nrxn1* heterozygous KO neurons (Cre) trans-differentiated using forced expression of Ngn2 from ES cells that were derived from heterozygous *Nrxn1* cKO mice (Chen et al., 2017).

Data are means ± SEM (n = cells/independent cultures are indicated in bars). Statistical analyses by Student's *t* test comparing test samples to controls uncovered no significant differences.

A

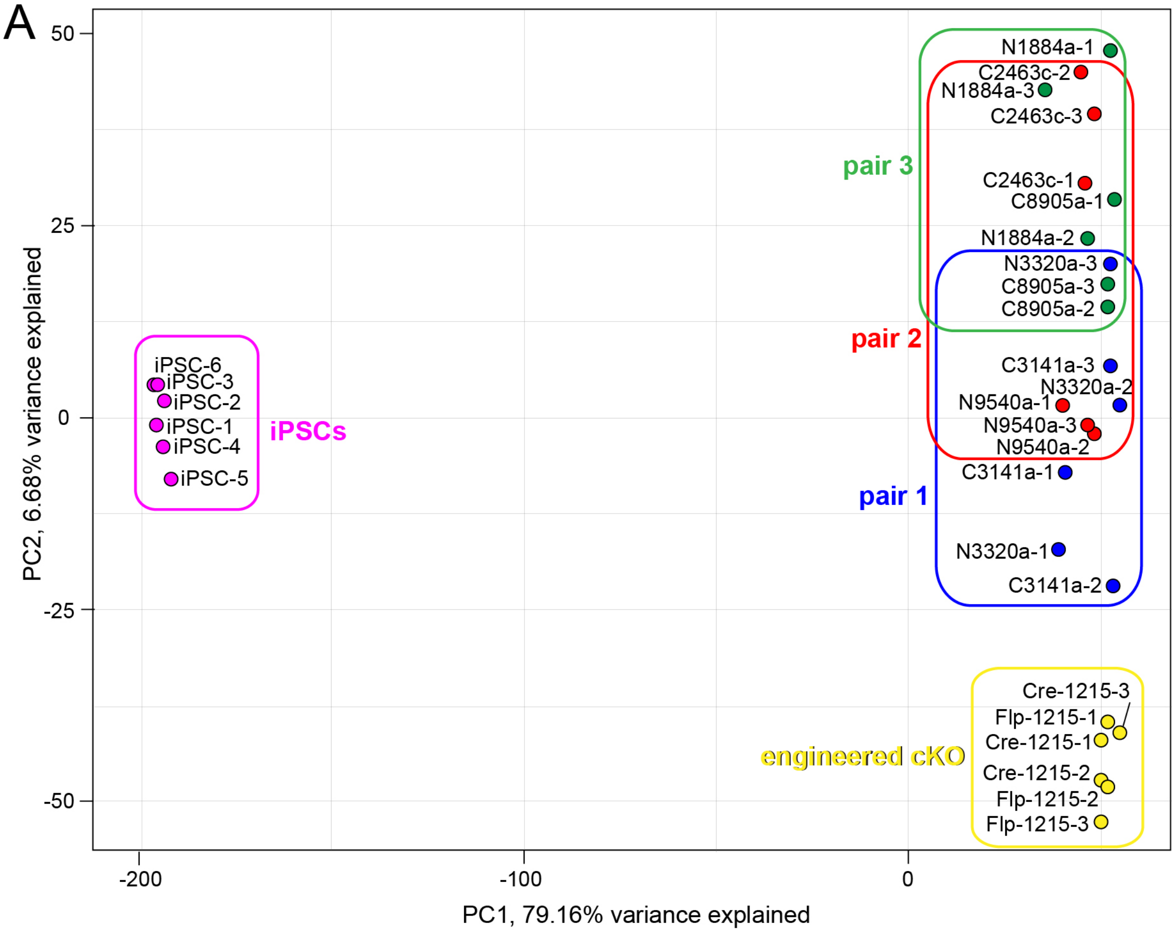


B

**Two-sample t-tests of patient-derived (*Nrxn1$^{del}$*) vs. control (Ctrl.) neurons**

| | GENO | N | Mean | SD | t | df | p-value |
|---|---|---|---|---|---|---|---|
| PC1 | Ctrl. | 9 | 48.99 | 3.97 | 0.8492 | 13.890 | 0.410 |
| | *Nrxn1$^{del}$* | 9 | 46.96 | 5.98 | | | |
| PC2 | Ctrl. | 9 | 16.96 | 21.90 | 0.3686 | 15.999 | 0.717 |
| | *Nrxn1$^{del}$* | 9 | 13.14 | 22.03 | | | |

Figure S6: **Principal Components Factor Analysis**

**A.** Plot of PC2 vs. PC1 for all 30 cultures. PCA was carried out in R using PCAtools (http://www.bioconductor.org/packages/release/bioc/html/PCAtools.html; Blighe K, Lun A (2020). PCAtools: PCAtools: Everything Principal Components Analysis. R package version 2.0.0, https://github.com/kevinblighe/PCAtools) using data for 15,000 post-QC genes for all 30 cultures. All iPS cell cultures are located to the left. The iN cultures are to the right, with all 6 engineered cultures at the bottom right. The heat map in Fig. 9B suggests there are large gene expression differences between iPS and iN cells, but also important differences in the iN cells generated from engineered conditional *NRXN1* cKO vs. non-engineered iPS cell lines. Here, PC1 (~80% of the variance) represents the iPSC-iN difference, and PC2 (~7 %) shows a gradient from engineered to non-engineered cells.

**B**. Two-sample t-tests showed that *NRXN1$^{del}$* and control cultures did not differ in their distribution of PC1 or PC2 scores.

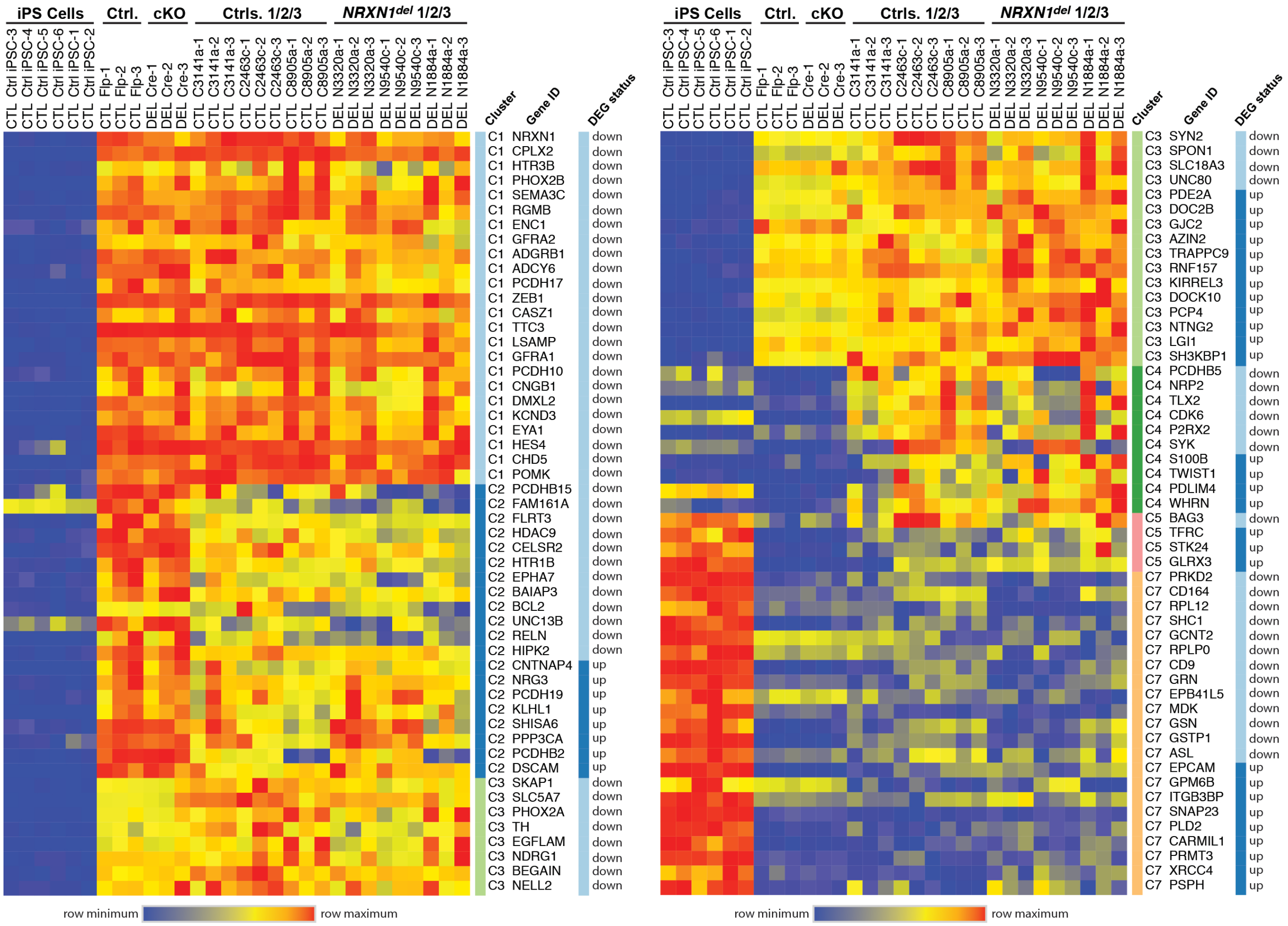


Figure S7: **Detailed view of the relative expression of specific differentially expressed genes (DEGs) in wild-type (WT) iPS cells and engineered and patient-derived heterozygous *NRXN1*$^{del}$ neurons (HET), and assignment of these DEGs to the clusters defined in Fig. 9**

Selected representative genes in each cluster are shown. DEGs are further categorized as up-regulated and down-regulated in comparisons of patient-derived *NRXN1*$^{del}$ vs. control neurons. For a complete list of genes in each cluster including genes that are unaffected by *NRXN1* heterozygosity, see Supplementary Data.

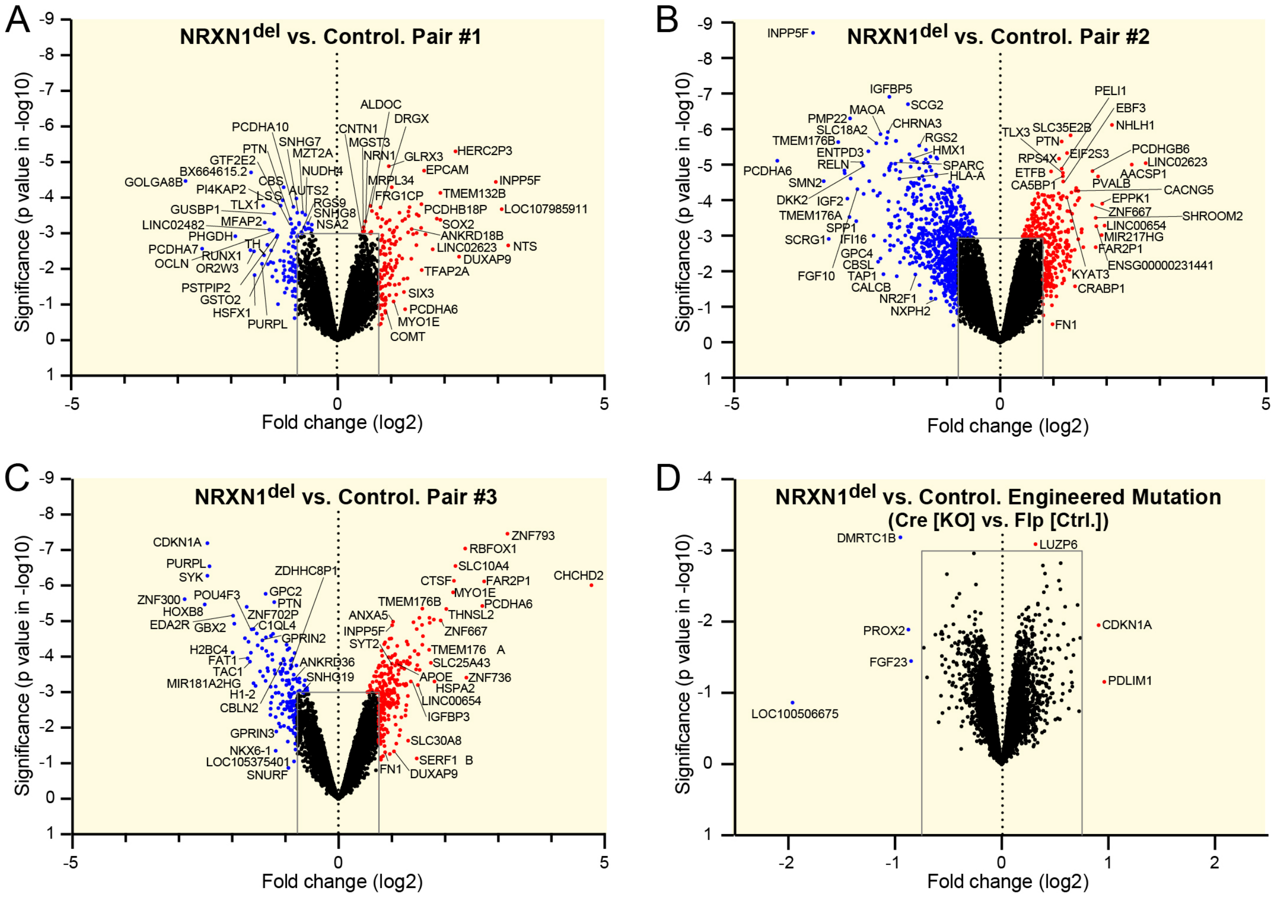


Figure S8: **Selected genes differentially regulated by *NRXN1*$^{del}$ in patient-derived and engineered induced neurons**

(**A-D**) Volcano plots of the DEGs identified by bulk RNAseq of neurons induced from pairs of control and patient-derived *NRXN1*$^{del}$ iPS cells (A-C) and from an engineered iPS cell line (1215) containing a conditional heterozygous *NRXN1* deletion (D). Significance (p-value in negative logarithmic scale) vs. fold change (in $\log_2$ scale) is plotted. Selected genes with the largest observed effects: |log2(fold-change)| ≥ 0.8 and/or -log10(P-value) ≥ 3 are highlighted on the plot. For full list of genes, see Supplementary Data.

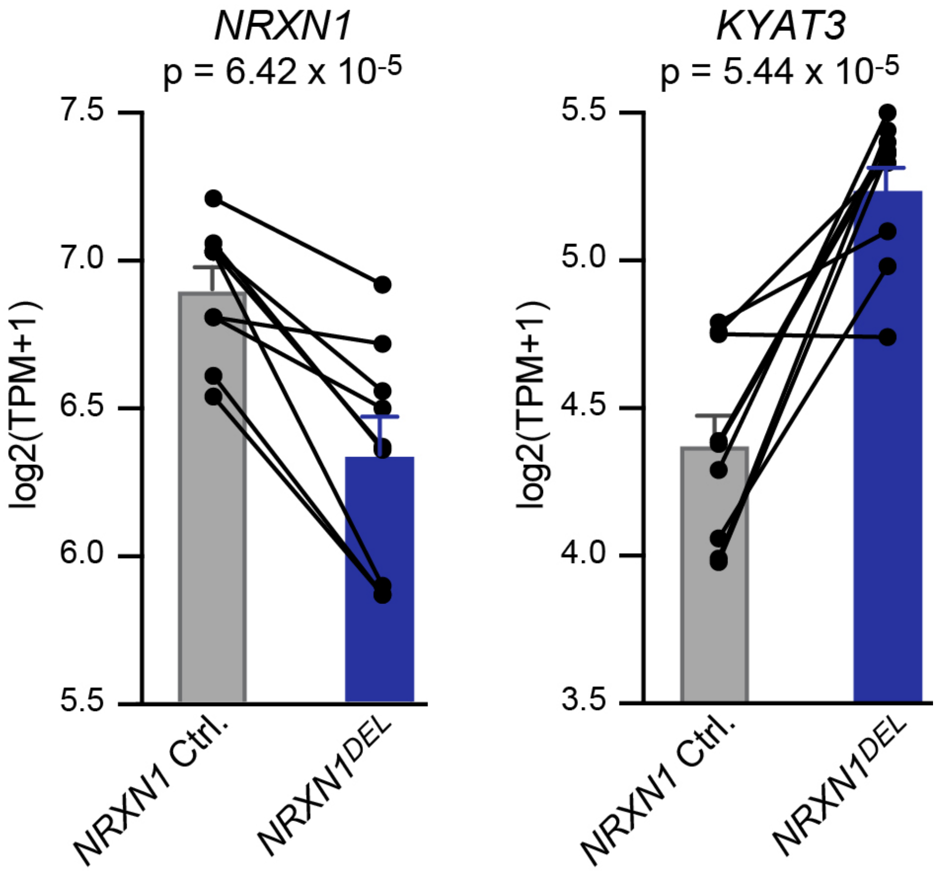


Figure S9: **Paired expression levels**

A moderated paired t-test (*limma-trend*) was used for all patient (deletion, DEL) vs. control (wild-type, WT) DEG tests, because specific pairs of patient and control replicate cultures were grown on the same plate at the same time to control for technical factors. Illustrated are log2(TPM+1) values for two DEGs (*NRXN1*, downregulated in patients, t=-5.953, p=6.42E-05, log2(fold-change)=-0.562; and *KYAT3*, upregulated in patients, t=6.060; p=5.44E-05; log2(fold-change)=0.87)), with lines connecting each replicate pair. Absolute expression values show variability across cultures (see Supplementary Data), but the slope of the pair difference is consistent for 9/9 pairs for *NRXN1* and 8/9 pairs for *KYAT3*.
